# Unusual molecular architecture of a human gut microbiota β-mannanase reveals a new CBM family

**DOI:** 10.64898/2026.02.11.704390

**Authors:** Natalia Łoś, Ieva Lelėnaitė, William G. T. Willats, Nicolas Terrapon, Ana Lorena Morales-García, Hamish C. L. Yau, Elisabeth C. Lowe, David N. Bolam

## Abstract

β-mannans are plant structural and storage polysaccharides prevalent in the human diet. Their degradation in the gastrointestinal tract is mediated by the human gut microbiota (HGM) through expression of a plethora of carbohydrate-active enzymes (CAZymes), although our understanding of the details of mannan breakdown is lacking. In this study, a prominent HGM member, *Bacteroides cellulosilyticus*, was found to be exceptionally efficient at utilising β-mannans, mediated by the expression of a single polysaccharide utilisation locus (PUL). Amongst the predicted surface CAZymes encoded in the PUL, we identified a family 26 glycoside hydrolase of an unusual molecular architecture. *Bc*WH2_GH26 contains a putative carbohydrate-binding module (CBM) directly intercalated into its catalytic domain, unlike classical CBMs which are located at the N- or C-termini of the catalytic domain. Phylogenetic and functional analyses of this internal CBM, and a homologue from another mannan user *Bacteroides uniformis*, revealed a narrow specificity for β-mannan and support their classification as a novel CBM family. To investigate the evolutionary basis for the unusual enzyme architecture, the effect of the CBM on the catalytic activity of the enzyme was assessed. No significant differences in the kinetic parameters were found between the full-length and CBM deletion constructs against both soluble and insoluble mannans. The potential role of the internal CBM in enzyme function is discussed in the context of the likely localisation of the *Bc*WH2_GH26 in the outer membrane utilisome encoded by the *Bc* mannan PUL.

## Introduction

Dietary fibre breakdown by the human gut microbiota plays a key role in host health through multiple mechanisms, including the production of beneficial metabolites like short chain fatty acids. Understanding the mechanisms by which fibre is broken down by the microbiota is thus key to our ability to modulate microbiota structure and composition for the benefit of health. β-mannans are a major source of dietary fibre as they are prominent components of many common foods, including legumes and seeds. In addition they are widely used in the food industry as additives such as thickeners and stabilisers(1). Despite this prevalence in the diet, to date, only limited number of studies focused on characterisation of β-mannan utilisation systems expressed by human gut microbes, such as *Bacteroides ovatus*(2) and *Roseburia intestinalis*(3).

Structurally, β-mannan in its simplest form consists of a β1,4-linked mannose backbone (homomannan) which can be further substituted with glucose (glucomannan), decorated with galactose side chains (galactomannan), or contain both types of modifications (galactoglucomannan)(1). The structural complexity of the polymer was found to vary between plant species and may depend on the plant’s developmental stage(4).

Glycoside hydrolases (GHs), the largest class of CAZymes, catalyse hydrolysis of glycosidic bonds within complex carbohydrates, and are to date grouped into nearly 200 families based on sequence similarities(5). GHs active on the β-mannan backbone belong to families 5, 26 and 113(5). GH26 is a well characterised family consisting primarily of endo-acting mannanases and exo-mannosidases. Many of the characterised bacterial GH26s, such as CtLIC26A (PDB: 2BV9), *B. ovatus* Man26A (PDB: 4ZXO), and *Bacillus subtilis* Man26 (PDB: 2WHK), consist of a single catalytic domain. In contrast, fungal GH26 mannanases often contain one or more carbohydrate-binding modules (CBMs)(6), often from family 35 at the N-or C-terminus of the catalytic domain, in common with most CBMs (7, 8).

To expand our understanding of mannan breakdown by the gut microbiota, we investigated the ability of a number of prominent human gut Bacteroidota species to utilise mannans. Two *Bacteroides spp., B. cellulosilyticus* and *B. uniformis* were particularly efficient mannan users and inspection of their predicted mannan-active surface CAZymes revealed GH26 enzymes with an unusual molecular architecture in that both enzymes contain a homologous domain of unknown function directly intercalated within their catalytic domains. Characterisation of this isolated domain revealed it to be a new family of mannan binding CBMs, hereafter named CBMXX. The phylogenetic distribution of the CBMXX as well as role of CBMXX in the GH26 enzymes indicate the unusual location of the CBM within the catalytic domain could be an adaptation to function within the surface mannan utilisomes of these species.

## Materials and methods

### Sources of carbohydrates

Mannooligosaccharides, cellooligosaccharides, carob galactomannan (low-viscosity), ivory nut mannan, konjac glucomannan (low-viscosity), wheat arabinoxylan, xyloglucan, barley and oat β-glucans were purchased from MegaZyme (Bray, Ireland). Guar gum galactomannan and hydroxyethyl cellulose were purchased from SigmaAldrich (MA, USA). Avicel was obtained from FMC Corporation (PA, USA). Insoluble mannan was obtained by washing 100 mg of ivory nut β-mannan in 1 ml deionised water five times and discarding the supernatant containing the soluble fraction of the polysaccharide. The pellet was then dried overnight at 37 °C.

### Growth array setup

Supplemented minimal medium (MM) was prepared at 2x concentration as per Table S1 and S2 with no carbohydrates added. β-mannan polysaccharides were prepared at 10 mg/ml and then mixed with the medium in 1:1 ratio, resulting in 1x MM and 5 mg/ml final carbohydrate concentration.

All growth arrays were conducted under anaerobic conditions, at 37°C. Nineteen different Bacteroidota species (Table S3.) were initially grown overnight (16-20 hours) in 5 ml of brain-heart infusion (BHI) broth (Sigma-Aldrich, MA, USA) with 1.2 µg/ml hematin and 0.2 mM histidine added.

Next, carbohydrate growth assays were conducted in triplicate in 96-wells plates. Each well contained 180 µl of 1x MM with 5 mg/ml selected carbohydrate, with an exception of three wells which did not contain any carbohydrate and served as no-carbohydrate negative controls. All the wells were then inoculated with 20 µl aliquot of the overnight culture. For the corresponding no-bacterium negative controls, 20 µl of BHI was added instead. Growth was monitored using Biotek Epoch Microplate Spectrophotometer (Agilent, CA, USA) at 15 minutes intervals over 48 hours.

No growth of any of the bacteria was observed in the no-carbohydrate wells. The mean of OD_600_ values obtained for the no-bacterium negative controls was subtracted from the growth values of corresponding strains. The obtained blanked growth values were then plotted as growth curves using GraphPad Prism 10.6.1. (MA, USA). Lastly, the maximum OD_600_ values reached during growth were used to create a heatmap.

### Phylogenetic analysis

To build the phylogenetic tree of the CBMXX family, 82 gene sequences were obtained from the CAZy (cazy.org)(5). The CBM domains were identified within each gene, extracted, and aligned with MAFFT using E-INS strategy. Next, BMGE (BLOSUM62 matrix) was used to select regions within the obtained multiple sequence alignment that were most suited for phylogenetic inference. The phylogenetic tree was inferred using IQ-tree webserver. Le-Gascuel (LG) was used as the substitution matrix, with the state frequency optimised by the Maximum-Likelihood (ML). Branch support was calculated using the ultrafast analysis of 1,000 bootstrap alignments.

### Cloning

*Bc*WH2_GH26 full-length construct without the predicted signal peptide referred to in the text either as *Bc*WH2_GH26 or *Bc*WH2_GH26 FL (SigP 6.0; amino acids 21-558), *Bu*_CBM (amino acids 240-354), and *Ruminococcus champanellensis* (RUM_21270; amino acids 32-616) were amplified by PCR from genomic DNA using KOD DNA Polymerase (Novagen) and primers presented in Table S4. The constructs were then cloned into pET-21a vector using NheI/XhoI, such that the recombinant protein encoded a C-terminal 6xHis-tag. To confirm correct insertion, the constructs were sequenced using T7 primers (Eurofins Genomics). The *Bc*WH2_CBM (amino acids 239-355) and the truncated *Bc*WH2_GH26_ΔCBM construct (amino acids 21-238, 356-558) were synthesised and inserted into a pET28a vector by Twist Bioscience with an N-terminal 6xHis-tag. The modularity of each construct is represented in Figure S1, and the full sequences are present in Table S5.

### Site-directed mutagenesis

The *Bc*WH2_CBM mutant constructs were synthesised and inserted into a pET-28a(+) vector by Twist Bioscience. In each case, a singular residue (W257, W301, Y310, H339) was substituted to alanine. The rest of the CBM sequence remained unchanged. The obtained constructs contained N-terminal and C-terminal 6xHis-tag. Full sequences are present in Table S5.

### Expression and purification of recombinant proteins

All the constructs were expressed using Tuner (DE3) competent *E. coli* cells. Cells containing the recombinant plasmid were grown in 1 L Luria-Bertani (LB) broth (containing either 50 µg/ml kanamycin or 100 µg/ml ampicillin) at 37 °C with shaking at 180 rpm to mid-exponential phase (OD_600_ ∼1.0). The recombinant protein expression was induced by adding 0.2 mM isopropyl β-D-1-thiogalactopyranoside. After incubating at 16 °C with 120 rpm agitation for 16 hours, the cells were harvested, and recombinant proteins were purified using immobilized metal ion affinity chromatography (IMAC). Cell-free extract was passed through 5 ml bed volume Talon™ resin column and washed with 20 ml of 20 mM Tris buffer pH 8.0 with 150 mM NaCl. The purified His-tagged proteins were eluted using 15 ml of 10 mM imidazole and 15 ml of 100 mM imidazole in Tris buffer. The proteins were then dialysed into and spin-concentrated using Vivaspin™ 10,000 MW concentrator columns (Cytiva, MA, USA) and the purity of the obtained proteins was assessed using SDS-PAGE gels stained with Coomassie blue (Abcam, Cambridge, UK).

### Enzyme assays

All the enzymatic reactions were carried out at 37 °C. To investigate the product profile of the *Bc*WH2_GH26 constructs, 0.5 μM final concentration of enzyme was incubated for various time points with 5 mg/ml mannan polysaccharides in 50 mM potassium phosphate buffer, pH 7.5. The reactions were stopped by boiling for 10 minutes and the products analysed using high-performance anion-exchange chromatography with pulsed amperometric detection (HPAEC-PAD; Thermo ICS-6000) using Dionex CarboPac PA300-4 µm column. Detection was facilitated by a gold working electrode and a PdH reference electrode, using a standard Carbo Quad waveform. Buffer A consisted of 100 mM NaOH, Buffer B was composed of 100 mM NaOH and 0.5 M sodium acetate, and Buffer D was a 500 mM NaOH solution. The following method was used: 0-55 min of linear gradient of 50% Buffer A; 55-60 min isocratic elution with 100% Buffer B; 60-65 min isocratic elution with 100% Buffer D; and lastly 65-75 min of linear gradient of 50% Buffer A.

To determine the specific activity and *k*_cat_/*K*_M_ values of the enzymes on soluble substrates, reducing sugar assays were carried out using 3,5-dinitrosalicylic acid (DNSA), as previously described(9). All reactions were carried out at 37 °C with constant 850 rpm agitation. Agitation was used to ensure adequate mixing of the reactions due to the viscosity of the mannans at the highest concentrations used. Each reaction had a final volume of 400 µl and contained 2.5, 3, 3.5, 4, 4.5 mg/ml of either CGM or KGM, and 50 mM potassium phosphate buffer, pH 7.5. The final enzyme concentration was 10 nM. At 5, 10, 15, 20 and 30 min timepoints, 60 µl aliquot was removed and the reactions were stopped by addition of DNSA reagent in a 1:1 volume ratio. The amount of reducing sugar released was quantified by measuring sample’s absorbance at 540 nm and using a standard curve of mannose. The *k*_cat_/*K*_M_ values were calculated from the initial slopes of the reactions as previously described(9), as the substrate concentrations used were significantly below the *K*_M_.

For the insoluble substrate, 3 mg/ml final concentration of insoluble ivory nut mannan was used with 1 µM final enzyme concentration in 50 mM potassium phosphate buffer, pH 7.5. The reactions were incubated at 37 °C with constant 1500 rpm agitation and timepoints were taken at 15, 30, 60, 120, 180, 240, 300, 360, 420 min, then every hour between 24-30 hours. One final timepoint was taken after 52 hours incubation. The reactions were stopped by adding DNSA reagent in a 1:1 volume ratio to the sample, as previously described(9). The concentration of reducing sugars was determined by measuring the *A*_540_ and comparing against a mannose standard curve. The data was then analysed using GraphPad Prism 10.6.1. (MA, USA).

### Affinity gels

Affinity gels were used for initial screening of the WT CBMs as well as initial analysis of the effects of the point mutations. Native polyacrylamide gels were made with 0.1% (w/v) final ligand concentration and 5 μg of each protein was loaded. Negative control gels did not contain ligand. Bovine serum albumin (BSA) (5 μg loaded) served as a negative control to exclude non-specific binding. Proteins were then visualised using Coomassie blue.

### Isothermal titration calorimetry

Isothermal titration calorimetry (ITC) using MicroCal PEAQ-ITC (Malvern Panalytical, Malvern, UK) was used to further investigate the specificity and affinity of the novel CBMs. Ligands were made up at either 5 mg/ml for polysaccharides or 5 mM for oligosaccharides in dialysis buffer (20 mM Tris, pH 8.0 with 150 mM NaCl). They were then injected in 18 x 2 μl aliquots into the sample cell containing 100 μM protein sample in the dialysis buffer to minimise heats of mixing. The titration was carried out at 25°C. The fitting of the isotherm to a single set of sites binding model was carried out using MicroCal PEAQ-ITC Analysis Software v1.41. The molar concentration of the available binding sites in the polysaccharide substrates was determined by fixing the value of n to 1. This assumption was deemed valid from the structural, mutagenesis and oligosaccharide binding data showing only a single binding site on the CBMs, in addition to close to sigmodal binding curves that allowed confident determination of stoichiometry from the fitted data.

### Insoluble polysaccharide binding assay

Insoluble polysaccharide binding was analysed using a pull-down assay as previously described(10), with some modifications. Briefly, 10 μg of purified CBM was incubated in 250 μl DI water, containing 4 mg insoluble ivory nut mannan (see ‘sources of carbohydrates’ above) or insoluble cellulose (Avicel), with 50 μg BSA to block non-specific binding. The negative control did not contain any ligand. The reactions were incubated for 30 min at 25 °C with 900 rpm agitation. The samples were then centrifuged at 10,000g for 15 min. Supernatant was collected and pellet was washed 5x in 20 mM Tris, pH 8.0 with 150 mM NaCl. After the final wash, 20 μl SDS loading buffer was added to the pellet. For the supernatant and negative control, 15 μl sample of each was taken and mixed with 5 μl SDS loading buffer. After boiling for 10 minutes, 12 μl of each sample was loaded on an SDS-PAGE gel. Proteins were visualised using Coomassie blue.

### Preparation of carbohydrate microarrays and microarray probing

Carbohydrate microarrays were prepared as previously described(11). Briefly, glycan standards were dissolved in deionised water to a final concentration of 2 mg/mL and placed on a rotary shaker at 4 °C for 16 h to ensure complete solubilisation. Glycan solutions were added to the wells of 384-well microtiter plate and diluted 1:1 (v/v) with printing buffer (47 % glycerol, 52.9 % deionised water, 0.06 % Triton X-100 and 0.04 % Proclin™ 200) and printed onto nitrocellulose membrane using a non-contact microarray robot (‘Marathon Argus’, Arrayjet, Roslin, UK).

Probing of printed glycoarrays was performed according to Johnsen et al. (2015) (12) with slight modifications. Briefly, microarrays were blocked for 1 h at room temperature in 1x PBS supplemented with 5 % (w/v) semi-skimmed milk powder (MP/PBS). Following incubation, arrays were probed for 2 h with the following probes: *Bc*WH2_CBM (His-tag); characterised β-mannan-binding CBM27A (His-tag, NZYtech)(13); anti-(1,4)-β-mannan BS-400-4 (mouse, BioSupplies); anti-heteromannan LM21 (rat) and anti-galactomannan CCRC-M75 (mouse, CarboSource) monoclonal antibodies. Monoclonal antibodies were used at 1:50 dilution, *Bc*WH2_CBM at 1:10 dilution, and CBM27A at 10 µg/ml. Microarrays were washed thoroughly in clean 1x PBS and then incubated for 2 h with alkaline phosphatase conjugated secondary antibodies (anti-His tag in the case of CBMs and anti-rat or anti-mouse in the case of mAbs) diluted 1:1000 in MP/PBS. mAb and binding was then detected by 5-bromo-4-chloro-3’-indolyphosphate p-toluidine (BCIP) salt and nitro-blue tetrazolium (NBT) chloride. Membranes were rinsed in fresh water and allowed to dry overnight. Microarray probing was performed in triplicate for each probe and control. Developed arrays were scanned at 2400 dpi and converted to negative JPEG files. Binding intensities of probes to individual spots was quantified using microarray analysis software (Array-Pro Analyzer Version 6.3). A value of 100 was assigned to the highest mean spot signal intensity recorded and all other values were normalised accordingly(14).

## Results

### Identification of β-mannan PULs in Bacteroides cellulosilyticus and Bacteroides uniformis

In pursuit of characterising novel β-mannan degradation mechanisms, we cultivated a total of 19 different human gut *Bacteroides* and *Phocaeicola* species using three types of β-mannan (carob galactomannan [CGM], konjac glucomannan [KGM], and guar gum galactomannan [GGM]) as a sole carbon source, Figure 1. Interestingly, despite both genera being known to be versatile glycan utilisers(15), we found that the majority of the tested species were unable to use all of the β-mannans for growth, suggesting a lack of β-mannan-specific enzymatic machinery, Figure 1A. In contrast, two of the species tested – *B. cellulosilyticus* and *B. uniformis* – grew exceptionally well on all three types of β-mannan used (shortest lag, fastest exponential and highest max OD; Figure 1B).

**Figure 1.**
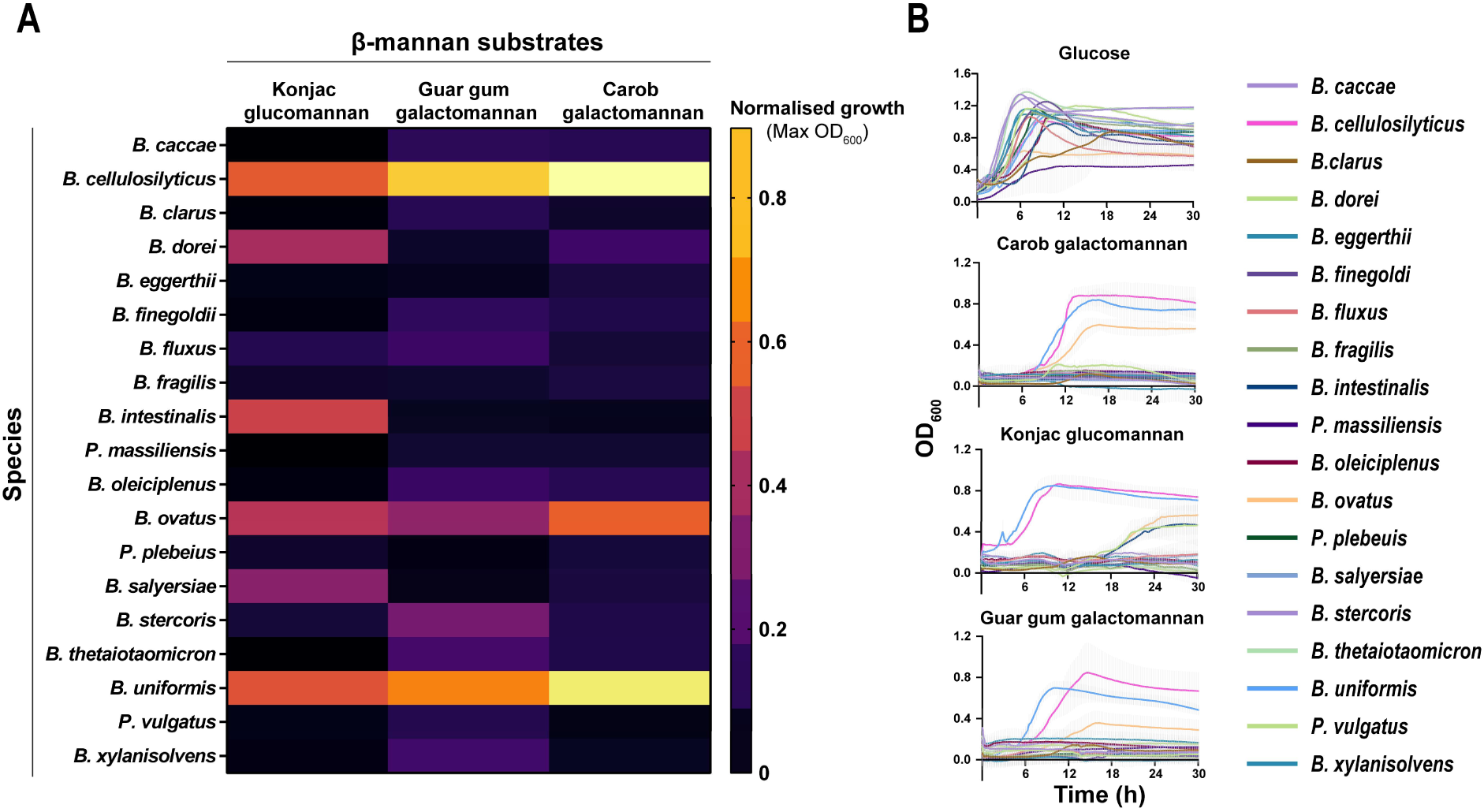
Growth of different gut Bacteroidota species on β-mannans. A) Heatmap showing maximum OD_600_ values reached by 19 human gut *Bacteroides* and *Phocaeicola* species when grown on 5 mg/ml β-mannan substrates – konjac glucomannan, guar gum galactomannan, and carob galactomannan. Bacteria were grown in 96-well plate (200 µl well volume) over a course of 48 h. The OD_600_ values of negative controls were subtracted before plotting. B) Corresponding growth curves used to plot the heatmap.

To investigate the mechanisms underlying these species ability to efficiently degrade mannans, we examined their genomes for potential mannan targeting enzymes. With *B. cellulosilyticus* WH2 a previous study has shown that PUL32 (spanning locus tags BcellWH2_02034-02020) were amongst the most highly upregulated during growth on β-mannan(16). This large and complex PUL is predicted to contain multiple mannan active CAZymes, including two GH26s (Table S6). Bioinformatic analysis then allowed us to identify a syntenic PUL in *B. uniformis* – Bu PUL9 (locus tags NQ510_02155-NQ510_02045). The two PULs share multiple homologous genes (highlighted in Figure 2), however, there are also key differences in their compositions. Notably, *Bc*WH2 PUL32 lacks multiple exo-glycosidases present in the *Bu* PUL9 (GH2, GH97, GH3/Peptidases), and the two PULs contain different phosphorylase families (GH130_1 in *Bc*WH2 PUL32 and GH94 in *Bu* PUL9). Despite these differences, the two PULs do share a prominent feature – they both contain homologous GH26 enzymes of an unusual molecular architecture. Furthermore, the two enzymes – *Bc*WH2_GH26 (locus tag: BcellWH2_02025) and *Bu*GH26 (NQ510_02065) – are predicted to be localised to the outer membrane (based on their type II signal peptide), suggesting a potential key role in the initial β-mannan chain breakdown. However, their most interesting feature is a previously uncharacterised domain of unknown function (DUF), inserted directly into the peptide sequence of their catalytic module, Figure 2. While the overall catalytic activity of *Bu*GH26 has previously been described(17), no functional characterisation of its internal domain was carried out. Both DUF domains have a β-sandwich-like fold with surface aromatics forming a potential binding cleft, an architecture common to many CBMs. Based on this we suspected the internal DUF of these enzymes could be a novel family of CBMs.

**Figure 2.**
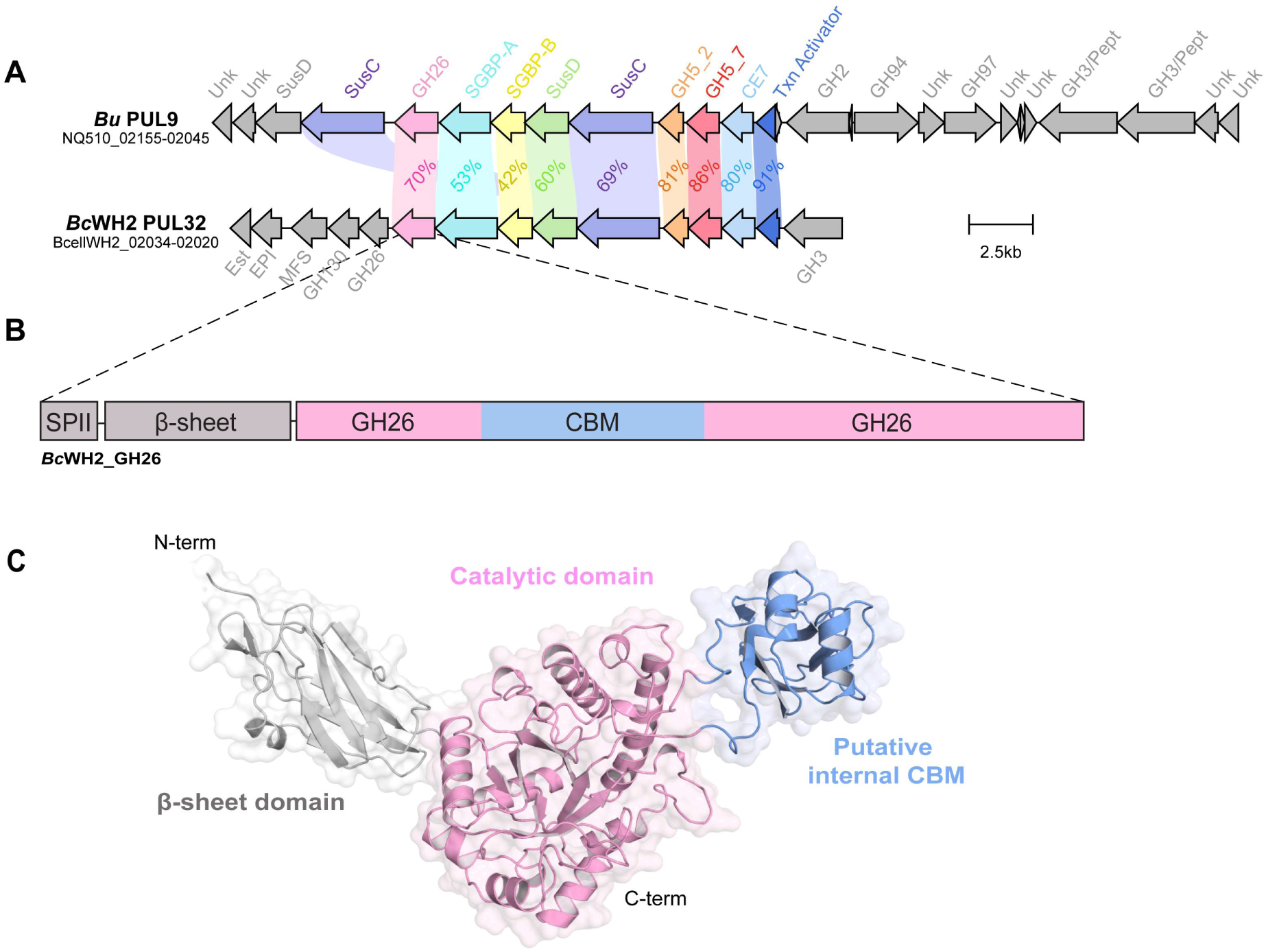
*B. cellulosilyticus* and *B. uniformis* β-mannan PULs and molecular architecture of *Bc*WH2_GH26. A) Map of *Bc*WH2 PUL32 and *Bu* PUL9, both containing GH26 enzymes with internal domains predicted to be CBMs. Composition of *Bc*WH32 PUL32 was determined experimentally, with all the genes shown to be upregulated during the bacterium’s growth on β-mannans (16). Composition of the syntenic *Bu* PUL9 was predicted using PULDB(18). To indicate similarities between the two PULs, genes sharing minimum of 30% amino acid sequence identity have been aligned and highlighted in different colours, with the % identity shown. The remaining non-homologous genes are shown in grey. Created and visualised using Clinker(19). B) Schematic representation of the molecular architecture of *Bc*WH2_GH26. The enzyme consists of a type II signal peptide (SPII, in grey), an N-terminal β-sheet domain of unknown function, hypothesised to be a spacer domain involved in positioning the enzyme in the surface utilisome (in grey), a catalytic GH26 domain (in pink), and a domain shown here to be an internal CBM (in blue). C) A model of the *Bc*WH2_GH26 structure, generated using AlphaFold2(20). Signal peptide was removed, and each domain was coloured as per schematic above*, i.e.,* β-sheet domain in grey, catalytic domain in pink, and CBM in blue.

### Internal DUFs of *Bc*WH2_GH26 and *Bu*_GH26 are highly specific β-mannan binding CBMs

Using carbohydrate microarrays, the putative internal CBM of *Bc*WH2_GH26 was initially screened against a broad range of polysaccharides selected as representatives of major glycan classes, and its binding specificity was compared to CBM27A (a known mannan binder) and well-characterised mannan-specific monoclonal antibodies (mAbs), BS-400-4 and LM21, 3A. This initial screen showed *Bc*WH2_CBM binding to CGM but not GGM. *Bc*WH2_CBM was more selective than CBM27A, which in addition to binding CGM, also showed signs of weak binding to several other glycans such as pectin and arabinoxylan, Figure 3A.

**Figure 3.**
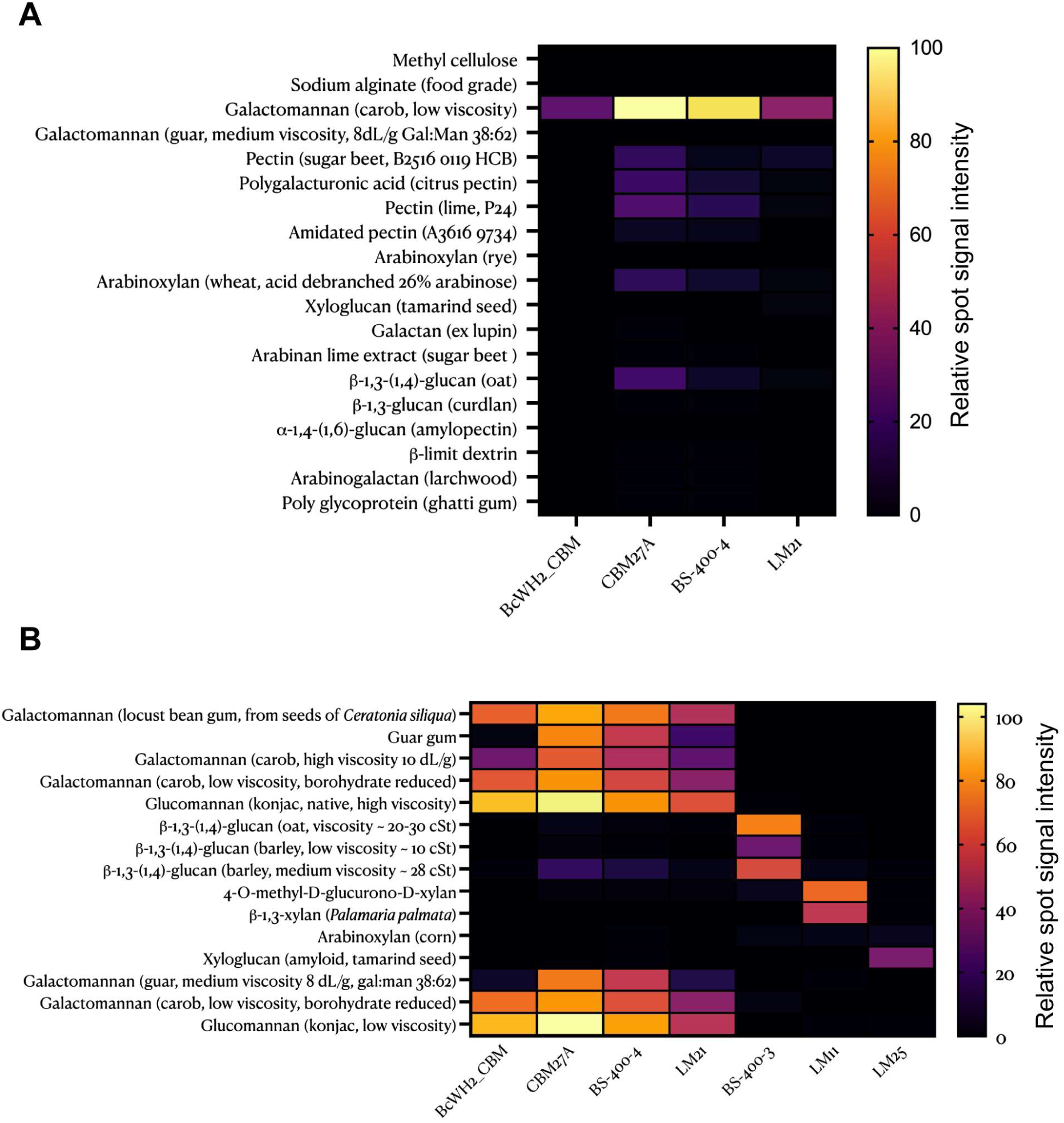
Binding specificity screening of *Bc*WH2_CBM using carbohydrate microarrays. Carbohydrate microarrays were used to obtain heatmaps showing results of broad-scope screening of *Bc*WH2_CBM to diverse polysaccharide standards (A) and more mannan-focused binding specificity screening (B). The polysaccharide standards are listed on the left. The mannan binding probes CBM27 and mAbs BS-400-4 and LM21 were used as positive controls, and mAbs BS-400-3, LM11 and LM25 were used as negative controls. Data are relative mean spot signals obtained from three independent printing and probing experiments and on each array, carbohydrate standards were printed as 3 technical replicates. The highest mean signal in the dataset was set to 100, and all other values were normalised accordingly.

Next, the binding specificity of *Bc*WH2_CBM was investigated in more detail using an extended set of β-mannan linked hemicellulose standards, Figure 3B. *Bc*WH2_CBM bound to all CGM preps tested but failed to bind to GGM, in contrast to CBM27A which bound to all types of galactomannan. *Bc*WH2_CBM also showed binding to KGM, but did not bind any glucans or xylans, further highlighting its narrow β-mannan specificity.

The results of the carbohydrate microarray screening were further confirmed by running native affinity gels. Both *Bc*WH2_CBM and its close homologue from *B. uniformis* GH26, *Bu*_CBM (∼70 % identity), bound CGM and KGM, but failed to bind to barley mixed-linkage β-glucan and wheat arabinoxylan, Figure S2.

### Assessing the affinity of *Bc*WH2_CBM and *Bu*_CBM

To further examine the specificity and affinity of these novel CBMs, isothermal titration calorimetry (ITC) was performed using a range of β-linked polysaccharides. *Bc*WH2_CBM and its homologue, *Bu*_CBM, were found to bind to both CGM and KGM, with comparable affinity, Table 1. Selected example ITC titrations are shown in Figure 4, with further ITC titrations shown in Figures S3 and S4 and full binding parameters set shown in Table S7.

**Figure 4.**
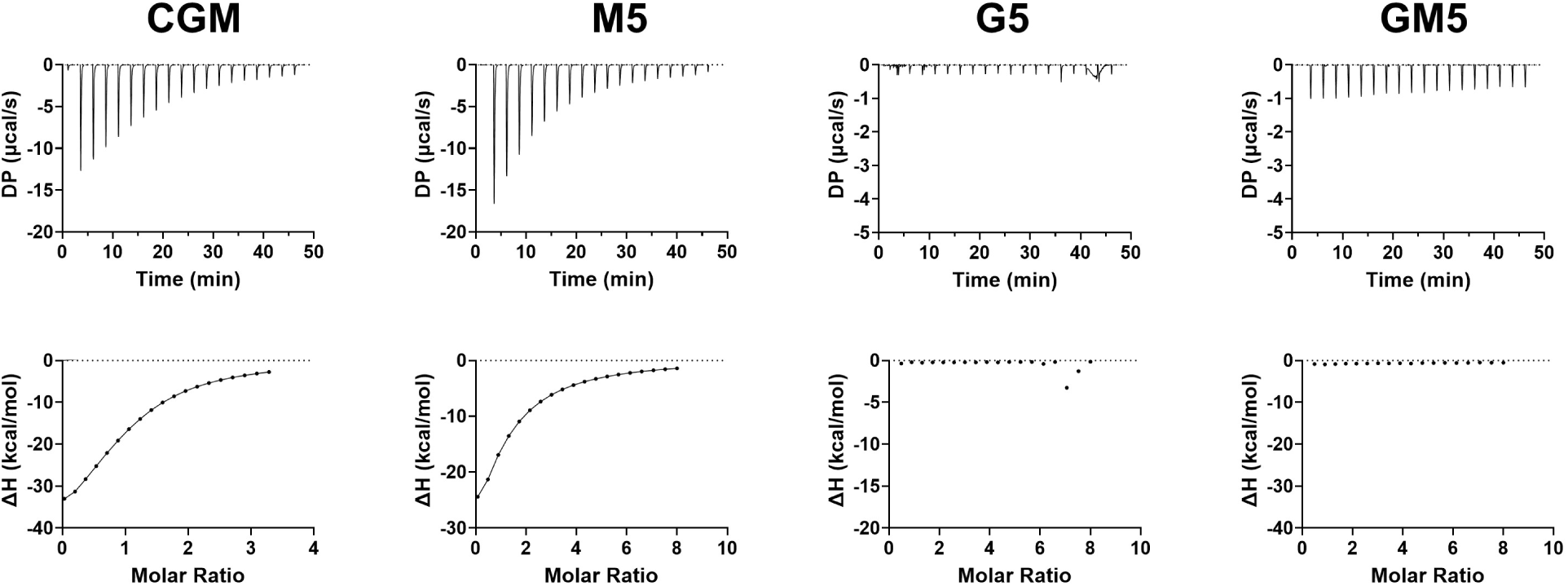
Example ITC of *Bc*WH2_CBM interactions with selected ligands. Glycans used are carob galactomannan (CGM), mannopentaose (M5), cellopentaose (G5), and di-alpha-D-galactosyl-mannopentaose (GM5). Titrations were conducted at 25 °C in 20 mM Tris buffer, pH 8.0, containing 150 mM NaCl. Upper parts of each panel are raw binding heats, lower parts are integrated data fit to single set of sites model if binding observed.

**Table 1.** Affinity of *Bc*WH2_CBM and *Bu*_CBM binding to oligo- and polysaccharides determined by ITC^a^.

| Construct | Ligand | $K_D$ (μM) ± SD | N ± SD |
| --- | --- | --- | --- |
| ***Bc*WH2_CBM<sup>b</sup>** | Carob galactomannan | 67 (± 6) | 1<sup>d</sup> |
| | Konjac glucomannan | 52 (± 7) | 1<sup>d</sup> |
| | Mannohexaose (M6) | 132 (± 34) | 1.3 (± 0.3) |
| | Mannopentaose (M5) | 235 (± 3) | 1 ± (0.3) |
| | Mannotetraose (M4) | 445 (± 21) | 1.7 (± 0.1) |
|  | Mannotriose (M3) | NB<sup>e</sup> | - |
|  | Hydroxyethylcellulose (HEC) | NB<sup>e</sup> | - |
| | Barley β-glucan | NB<sup>e</sup> | - |
|  | Xyloglucan | NB<sup>e</sup> | - |
|  | Cellopentaose | NB<sup>e</sup> | - |
| | 6<sup>3</sup>,6<sup>4</sup>-di- α-D-galactosyl-mannopentaose | NB<sup>e</sup> | - |
| ***Bu*_CBM<sup>c</sup>** | Carob galactomannan | 38 (± 6) | 1<sup>d</sup> |
| | Konjac glucomannan | 47 (± 21) | 1<sup>d</sup> |
| | Mannohexaose (M6) | 43 (± 2) | 0.8 (± 0.1) |
| | Mannopentaose(M5) | 82 (± 6) | 1.3 (± 0.4) |
| | Mannotetraose (M4) | 335 (± 26) | 0.7 (± 0.4) |
|  | Mannotriose (M3) | NB<sup>e</sup> | - |
|  | Hydroxyethylcellulose (HEC) | NB<sup>e</sup> | - |
|  | Xyloglucan | NB<sup>e</sup> | - |
|  | Cellopentaose | NB<sup>e</sup> | - |
| | 6<sup>3</sup>,6<sup>4</sup>-di- α-D-galactosyl-mannopentaose | NB<sup>e</sup> | - |

<sup>a</sup> Parameters determined by ITC. Standard deviation calculated for the means of at least triplicate titrations.
<sup>b</sup> *Bc*WH2\_GH26 isolated CBM.
<sup>c</sup> *Bu*-GH26 isolated CBM.
<sup>d</sup> N was set at 1 for polysaccharides (see Methods).
<sup>e</sup> No binding detected.

ITC with defined oligosaccharides revealed the minimal chain length required for binding was four mannoses, as the modules failed to bind to mannotriose. The chain length of the ligand did substantially influence the modules binding affinity, with *Bc*WH2_CBM affinity for M5 (*K*_D_ = 235 µM) decreasing two-fold compared to M6 (*K*_D_ = 132 µM) and further two-fold for M4 (*K*_D_ = 438 µM), suggesting the full binding site spanned at least 6 mannoses, although the number of binding sites on the CBM (N) for all manno-oligosaccharides was close to 1 for both CBMs, Table 1. Interestingly despite binding glucomannan, neither CBM bound any of the β-glucan homopolymers tested or cellopentaose. Furthermore, both while both CBMs could bind galactomannan, they were unable to accommodate di-galactosyl side chains on neighbouring mannose residues, as shown by lack of binding to 6^3^,6^4^-di- α-D-galactosyl-mannopentaose, despite binding M5. *Bc*WH2_CBM preference for less-decorated polysaccharides was also confirmed via native affinity gels, where a higher degree of retardation was observed for carob galactomannan (4:1 Man:Gal) compared to guar gum galactomannan (Man:Gal 2:1), Figure S2.

### *Bc*WH2_CBM is able to bind insoluble β-mannan

One of the classical roles of CBMs is thought to be to potentiate the activity of the cognate catalytic domain against insoluble substrates(21). To investigate the ability of *Bc*WH2_CBM to bind insoluble polysaccharides we carried out pulldown assays with the insoluble fraction of ivory nut β-mannan (INM) and microcrystalline cellulose (Avicel). Bovine serum albumin (BSA) was added to each reaction to minimise the effects of non-specific binding. The results revealed that *Bc*WH2_CBM is able to bind insoluble β-mannan but not cellulose, as evidenced by a band on the gel in the pellet fraction of mannan, but not Avicel, Figure 5. However, a band was also seen in the corresponding supernatant fraction vs mannan, suggesting that all available binding sites on the mannan have been saturated with CBM. The lack of binding to cellulose observed further confirmed the CBMs tight specificity for mannan.

**Figure 5.**
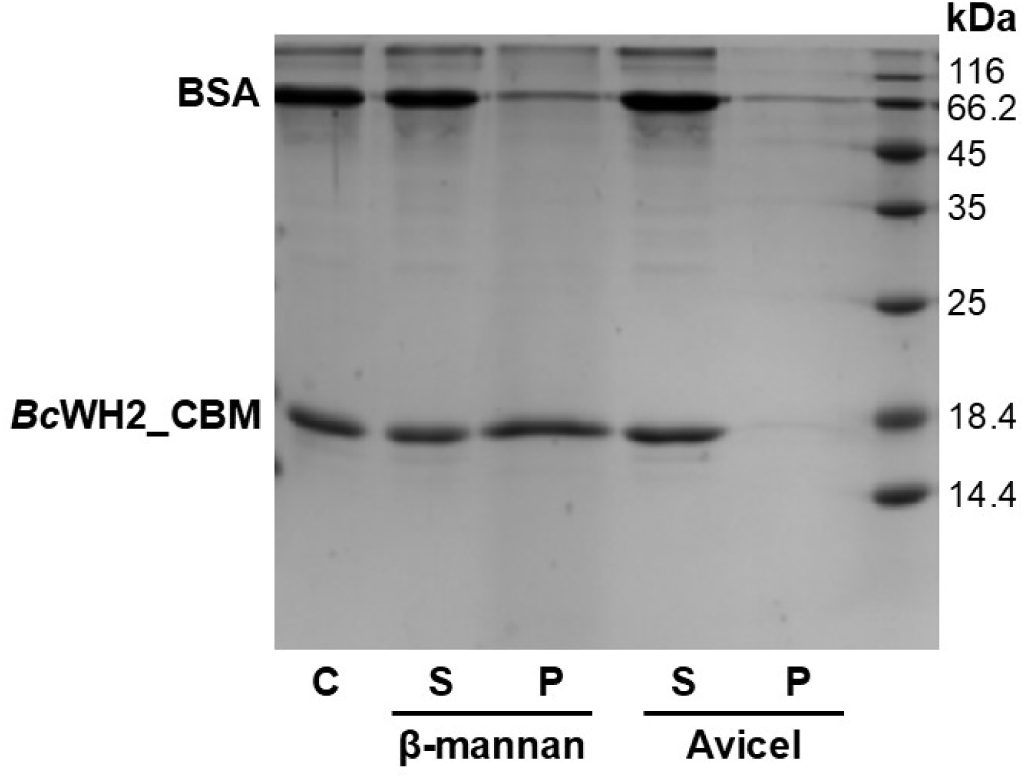
Binding *Bc*WH2_CBM to insoluble β-mannan. After incubating *Bc*WH2_CBM with insoluble mannan or cellulose (Avicel), the samples were centrifuged to separate supernatant (S) and pellet (P) fractions. The control (C) sample did not contain any polysaccharide. Bovine serum albumin (BSA) was included in each assay to minimise non-specific binding and as a non-interacting control. All samples were subsequently analysed by SDS-PAGE. Molecular weight standards are shown on the right.

**Figure 6.**
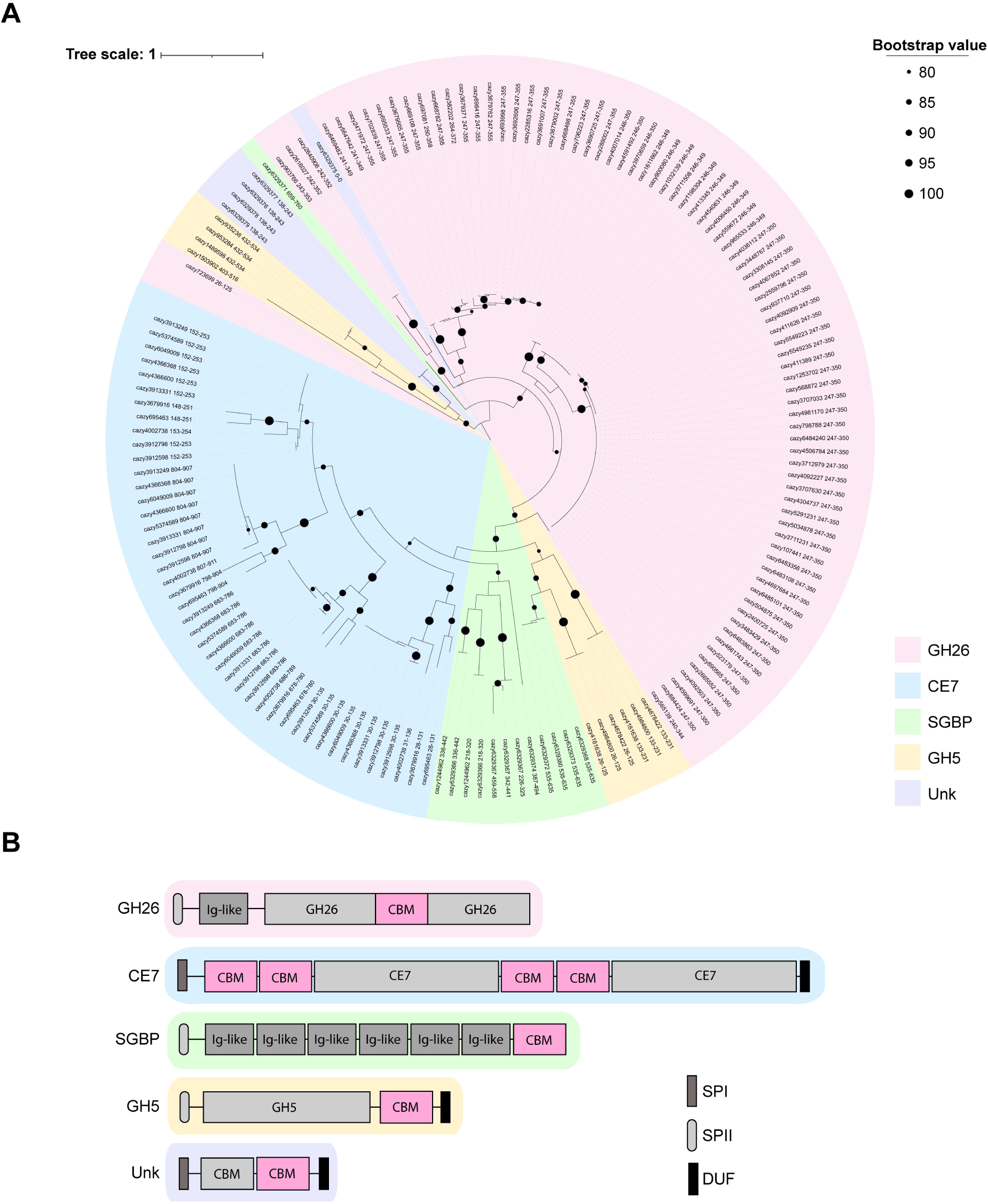
Phylogenetic analysis of the CBMXX family. A) Phylogenetic tree of all members of the CBMXX family. Sequences obtained from the CAZy database(5) and annotated with their CAZy database entry names. Each protein class associated with the CBMXX is indicated by a different colour, as follows: GH26 in pink, CE7 in blue, putative surface glycan-binding proteins (SGBP) in green (function hypothesized based on their sequence and structural similarity to previously characterised SGBPs), GH5 in yellow, and proteins of unknown function in purple. Nodes with bootstrap values greater than 80 are indicated by black circles of increasing size, as per legend. B) Schematic of domain architecture of representatives of each protein class associated with CBMXX. CBMXX is highlighted in pink, the remaining domains are depicted in shades of grey/black. Type of a signal peptide (SPI/SPII) and predicted protein functions have been annotated (GH26, CE7, GH5, CBM). Ig-like β-sheet domains present in putative SGBPs and GH26s are depicted as grey boxes. Additionally, CE7, GH5, and proteins annotated here as Unknown (Unk), all contain a C-terminal domain of around 30-40 amino acids of unknown function. These C-terminal modules are annotated here as domains of unknown function (DUF) and depicted as black rectangles.

### *Bc*WH2_GH26 CBM and *Bu*_CBM are founding members of a novel CBM family

Upon confirming *Bc*WH2_CBM and *Bu*_CBM to be novel β-mannan binding CBMs, a phylogenetic analysis was performed. This allowed for the establishment of a novel CBM family, CBMXX. Searching CAZy and PULDB databases (5, 18), we found 82 proteins to contain the CBMXX modules, Figure 1A. The majority of these were Bacteroidota proteins, found in the human and mammalian gut and oral cavity. CBMXX modules can be appended to various protein classes. Most often, they are intercalated into the catalytic domain of GH26 modules of *Bacteroidales* spp (*Bacteroides*, *Prevotella*, and *Duncaniella*) analogously to *Bc*WH2_GH26. In rare cases, such as in GH26 from *Prevotella sp.* AGR2160 (RefSeq. WP_081657628.1) the catalytic domain remains uninterrupted and is instead preceded by the CBM, indicating the CBM is not absolutely required to be located internal to the catalytic domain for functionality. CBMXX is also found associated with esterases, such as CE7 from *Segatella copri* (WP_317576045.1). In this case, two repeats of the CBM domain are followed by a CE7 module and this structure then repeats, Figure 1B. Less commonly, CBMXX can be appended at the C-term to a GH5. Unlike the CBMXX containing GH26s, these GH5s do not possess an extra Ig-like N-term domain. Interestingly, CBMXX are also found in putative surface glycan-binding proteins (SGBPs), based on the protein molecular architecture (multiple Ig-fold domains followed by the CBM at the C-term) and position of the gene within the PUL downstream of the *susD*. Lastly, in rare cases, members of this family can be found as discrete entities alongside another domain of a predicted structure suggesting a CBM of an unknown family. These double CBM proteins contain type I signal peptide, type (*e.g.,* GTC17253_17770), suggesting a different role to the other family members. Overall, this analysis suggests that CBMXX’s function may not be solely limited to the classical model of the CBM assisting the catalytic activity of enzyme. Aiming to investigate this further, we then looked at a broader genomic context of these CBMs.

### CBMXX is most often found within predicted β-mannan PULs

To examine the broader context in which CBMXXs function, we carried out genomic analysis. This revealed that 80-94% of the CBMXX family members are found within a CAZyme/PUL context. Out of the 82 proteins containing CBMXX module, 64 are located within classical PULs, and 2 are associated within CAZyme clusters. Furthermore, 11 out of the 16 remaining members are found within the neighbourhood of CAZyme genes such as GH26 and GH130, however not in a close enough proximity to be considered a cluster. Interestingly, four of the CBMXXs not found in the context of a PUL are the aforementioned discrete CBMs, not attached to any other protein class.

Focusing on the classical PULs, we found that although their exact composition varies, some patterns can be discerned. The GH family most often observed to co-occur alongside CBMXX are GH26, which were found in 91% of PULs containing the CBMXX, Figure 7. This was followed by GH5_7 present in 78% of CBMXX PULs, and GH5_2 and GH3, each found in 59% of PULs. Other co-occurring enzymes belong to families GH2 (47%), GH94 (42%), GH130_1 (36%) and GH97_3 (36%). GH26 and GH5_7 families predominantly consist of β-mannanases, meanwhile GH5_2 and GH3 are often involved in β-glucan degradation(5). Based on these previously observed family activities, our data suggest that CBMXXs are most often found in β-mannan or β-glucomannan targeting PULs. The latter is further supported by the presence of GH130_1 enzyme in over a third of CBMXX-containing PULs, as this GH130 subfamily contains solely mannosylglucose phosphorylases(5).

**Figure 7.**
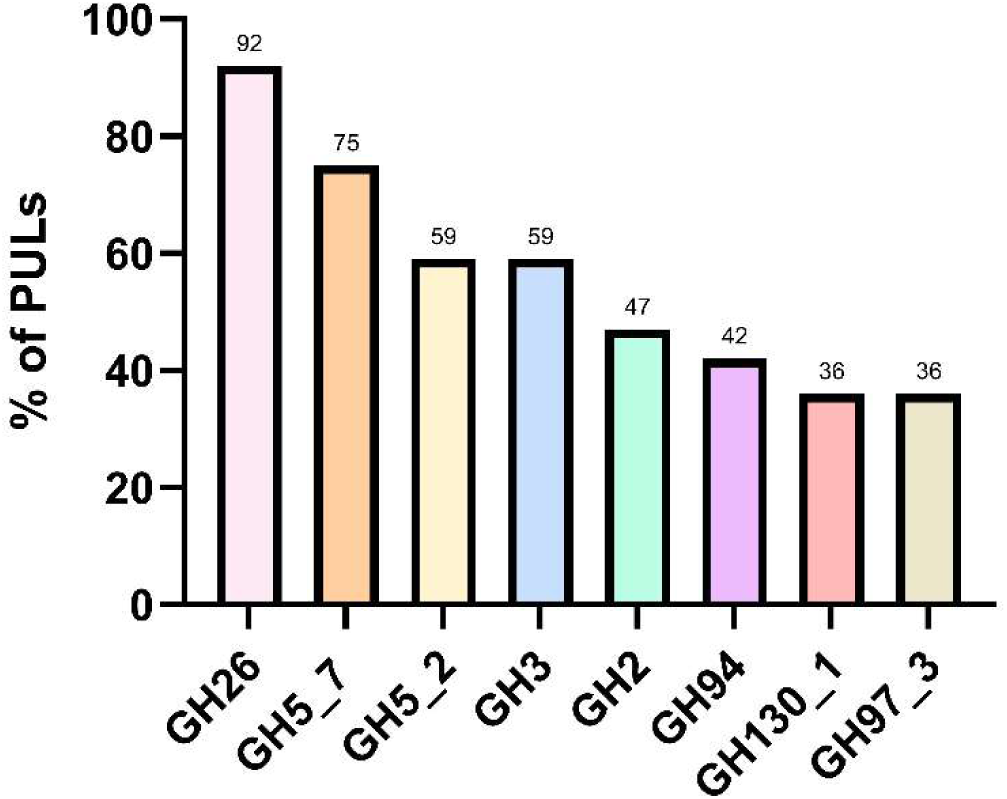
Co-occurrence of different GH families in PULs containing CBMXX. Depicted here as percentage of occurrence of a given GH family in all PULs in PUL-DB containing CBMXX(**18**). Data show that GH26 is a family most often associated with CBMXX PULs, as they can be found in 91% of these PULs. Other enzymes most commonly found within these PULs belong to families GH5_7 (found in 75% of CBMXX PULs), GH5_2 (59%), GH3 (59%), GH2 (47%), GH94 (42%), and GH130_1 and GH97_3 (36% each). Based on the activities previously observed in these families, these data suggest that CBMXXs are most often found in putative β-mannan or β-glucomannan targeting PULs.

Interestingly, our analysis did not reveal GH36 to be associated with the CBMXX PULs, even though this family of α-galactosidases is often involved in debranching galactomannans (22). Therefore, we speculate that either the specificity of the CBMXX PULs may lean more towards linear mannan/glucomannan utilisation, as opposed to galactomannan; or these PULs might instead utilise GH97 to remove α-galactose side chains, although this family has not previously been shown to act on β-mannan. Alternatively, the PUL enzymes might cooperate with a GH36 located outside the PUL. This may be the case for *B. cellulosilyticus*, where an orphan GH36 is upregulated during the growth on β-mannan at the same time as PUL32 containing CBMXX(16).

### Determining key ligand binding residues in *Bc*WH2_CBM

The putative ligand binding site of the *Bc*WH2_CBM was predicted based on the AlphaFold2 prediction of the protein structure. The degree of conservation of residues within the binding cleft varies, however, the two surface exposed tryptophan residues (W257 and W301) are conserved across the whole CBMXX family, suggesting their key role in ligand binding, Figure 8A. To experimentally confirm this, we made alanine mutants of four *Bc*WH2_CBM residues located within the putative binding site - W257, W301, Y310 and H339, Figure 8B.

**Figure 8.**
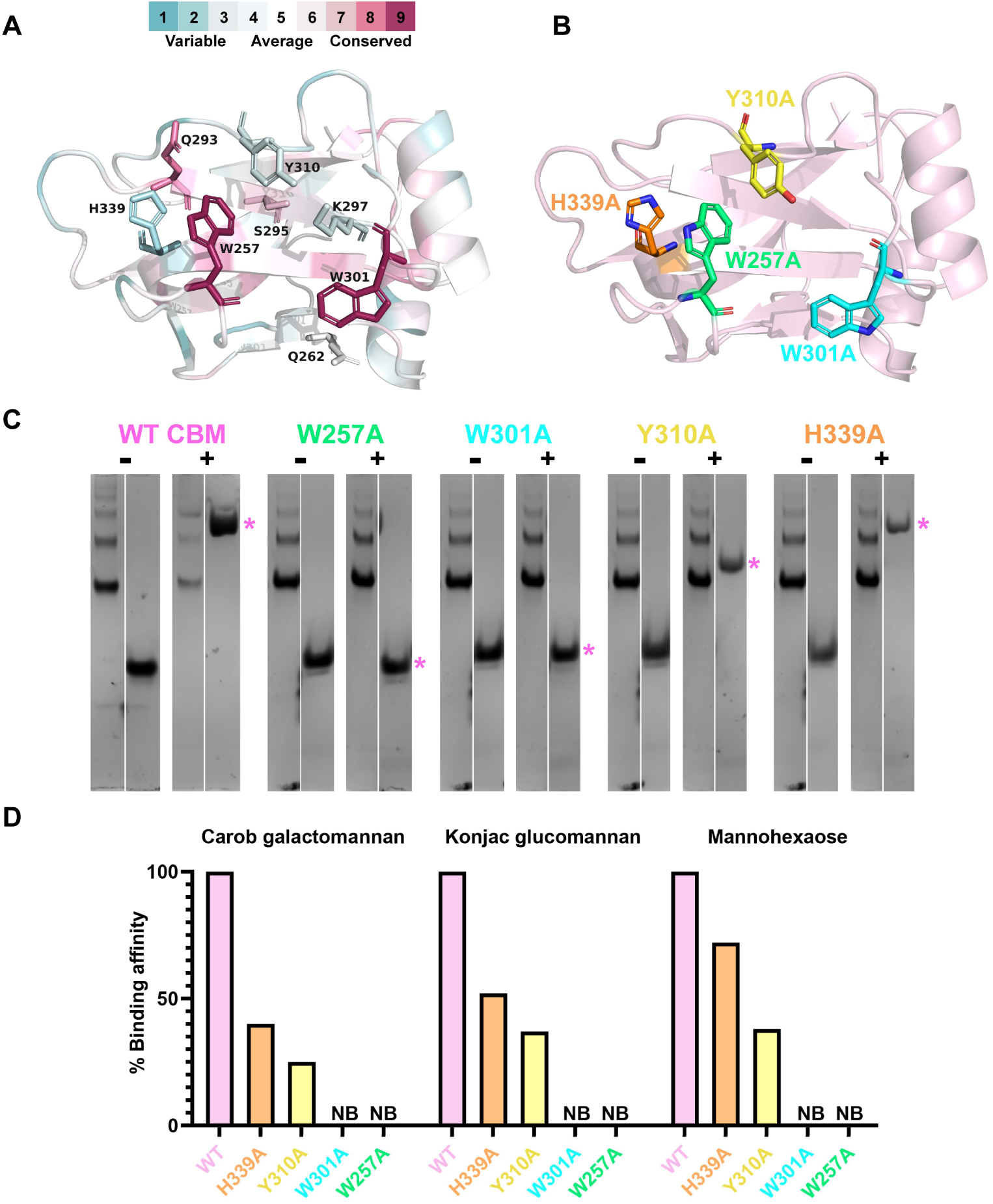
Identification of *Bc*WH2_CBM ligand binding residues. A) Conservation of proposed binding site residues created using ConSurf(23). Multiple-sequence alignment of the CBMXX family was used as a template. Conservation grades scale is shown. B) Highlighted residues within the proposed *Bc*WH2_CBM binding site, selected for mutagenesis C) Native gel containing 0.1% (w/v) konjac glucomannan (+) or no ligand (-) showing affinity of wild-type *Bc*WH2_CBM (WT) and its corresponding mutants to this ligand. BSA was used as a negative control and is shown on the left of each gel. The positioning of the CBM in each ligand-containing gel is indicated by a pink asterisk. D) *K*_D_ values of WT *Bc*WH2_CBM and its corresponding mutants for three ligands, shown here as percentage of the WT *K*_D_ value. NB indicated no binding was detected.

Affinity gel electrophoresis with glucomannan revealed that W257A and W301A mutants displayed no binding to the polysaccharide, meanwhile Y310A retained limited binding capacity and the H339A mutant bound similarly to WT on the gels, Figure 7C and S5.

To obtain more specificity and affinity data, the binding of the mutants to a range of oligo- and polysaccharides was assessed via ITC and compared against the wild-type protein, Figure 8D and Table S8. Once again, no binding was observed for W257A and W301A mutants on any of the ligands tested. The H339A mutant was able to retain almost half of the WT CBM’s binding affinity to the polysaccharides tested, however, interestingly, the mutation had less of an effect when protein was assayed against mannohexaose. This may be explained by the location of the H339 residue within the binding site and the differences in length between the ligands tested. Meanwhile, Y310A mutation decreased the ability of the CBM to bind its ligands, including mannohexaose, almost four-fold, Figure 7D.

Overall, these data confirm W257 and W301 to be key binding residues. Without the stacking interactions provided by them, the CBM completely fails to bind to its ligands. Y310 was found to also be important in ligand binding, however, to a lower degree than the two tryptophan residues. Due to its orientation within the binding site, we suspect that Y310 does not participate in stacking interactions, and instead, is involved in hydrogen bonding with the sugar. Similarly, H339 is most likely involved in H-bonding with ligand, albeit, to a lesser extent.

### The effect of the CBM on the enzyme’s activity vs soluble and insoluble mannans

To investigate the effect of the CBM on the activity and product profile of *Bc*WH2_GH26, a truncated construct of the enzyme was created, lacking the internal CBM (*Bc*WH2_GH26_ΔCBM). To ensure the stability of this construct, we performed nanoDSF scan of both proteins and did not observe significant differences in their melting temperatures, Figure S6.

The full-length enzyme (*Bc*WH2_GH26 FL) and *Bc*WH2_GH26_ΔCBM were incubated overnight with either CGM, KGM or M6, and the reaction products were analysed via HPAEC-PAD. No differences in the oligosaccharide products released by the two constructs were seen, Figure S7. Both enzyme variants released a mixture of oligosaccharides of various degree of polymerisation (DP), with the main products ranging in length from DP2 to DP6. The retention time of the oligosaccharides obtained from CGM and KGM hydrolysis differed compared to the linear standards, suggesting that they were either GlcMan heterooligosaccharides (for KGM) or branched oligosaccharides (for CGM), however, their exact identity has not been elucidated. Overall, the obtained data indicate an endo-acting mode of action of *Bc*WH2_GH26, as previously observed in enzyme’s homologs such as *Buni*_GH26(17) and GH26B from *Bacteroides ovatus*(22).

Next, the catalytic efficiency (*k*_cat_/*K*_M_) was determined for both the full-length protein and its truncated version against soluble substrates – CGM and KGM. Overall, both constructs were ∼1.5-2-fold more active against KGM compared to CGM, Table 2 **Table 3**. No significant differences were seen when comparing the two constructs on either CGM or KGM (paired t test p value > 0.05), further evidencing that the CBM does not increase the efficiency of the enzyme against soluble substrates, which agrees with a classical role of a CBM aiding efficiency against insoluble polysaccharides but being superfluous against soluble substrates(24, 25). The linear regressions used to estimate *k*_cat_/*K*_M_ are shown in Figure S8.

**Table 2.**
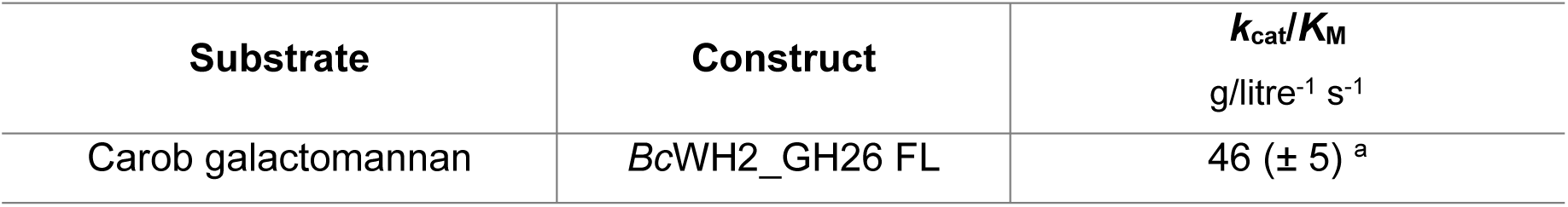
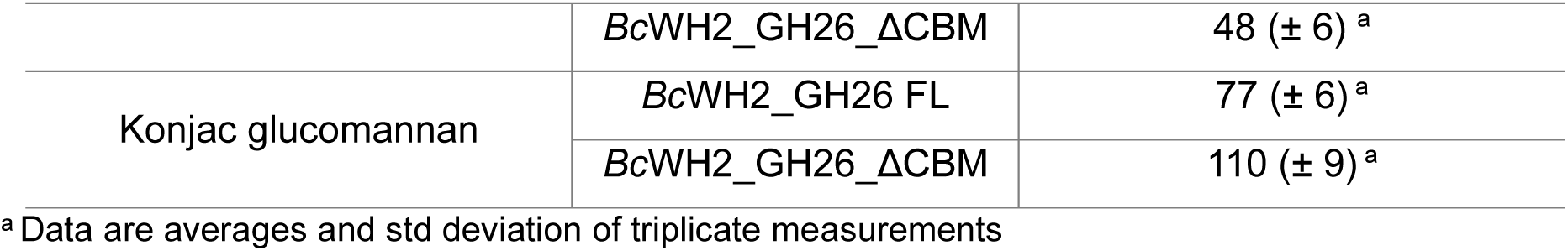
Catalytic efficiency of *Bc*WH2_GH26 FL and *Bc*WH2_GH26_ΔCBM against soluble mannans.

| Substrate | Construct | $k_{cat}/K_M$<br>g/litre<sup>-1</sup> s<sup>-1</sup> |
| --- | --- | --- |
| Carob galactomannan | *BcWH2_GH26 FL* | 46 (± 5)<sup>a</sup> |
|  | *BcWH2_GH26_ΔCBM* | 48 (± 6) <sup>a</sup> |
| Konjac glucomannan | *BcWH2_GH26 FL* | 77 (± 6) <sup>a</sup> |
|  | *BcWH2_GH26_ΔCBM* | 110 (± 9) <sup>a</sup> |

<sup>a</sup> Data are averages and std deviation of triplicate measurements

To test the effect of the CBM on insoluble substrate, specific activity of the two *Bc*WH2_GH26 constructs was determined using a reducing sugar assay with 3 mg/ml final concentration of the insoluble fraction of the ivory nut β-mannan (INM). The results were then compared against those obtained for equivalent concentration of soluble substrates (CGM and KGM), revealing, as would be expected, higher hydrolysis rates against soluble substrates compared to insoluble INM, Table 3. The rates of the two constructs were near identical at the initial stages of the reactions, Figure S9, resulting in only marginal differences in the calculated specific activity against all three substrates, including INM, Table 3. Notably though, after 48 hours we observed complete solubilisation (clearing) of the insoluble mannan by the *Bc*WH2_GH26 FL, whereas *Bc*WH2_GH26_ΔCBM variant failed to solubilise the mannan to the same extent, Figure S10A. Upon further testing, involving assaying a sample of the reaction against soluble CGM, we found that this was due to the differences in stability between the two constructs, with *Bc*WH2_GH26_ΔCBM having lost activity after the long incubation time of the assay, Figure S10B. To rule out stability issues affecting the solubilisation phenomenon observed, we repeated the initial assay with additional fresh aliquots of the enzyme added at 7 and 24 h incubation timepoints. This showed that *Bc*WH2_GH26_ΔCBM is also able to completely solubilise INM, Figure S10C, indicating that the internal CBM is not essential for this process. Furthermore, we found that this phenomenon is not a property unique to *Bc*WH2_GH26, as *Ruminococcus champanellensis* GH26 (*Rc*GH26; RUM_21270) containing an N-term CBM, was also able to solubilise INM in the same manner, Figure S10D.

**Table 3.**
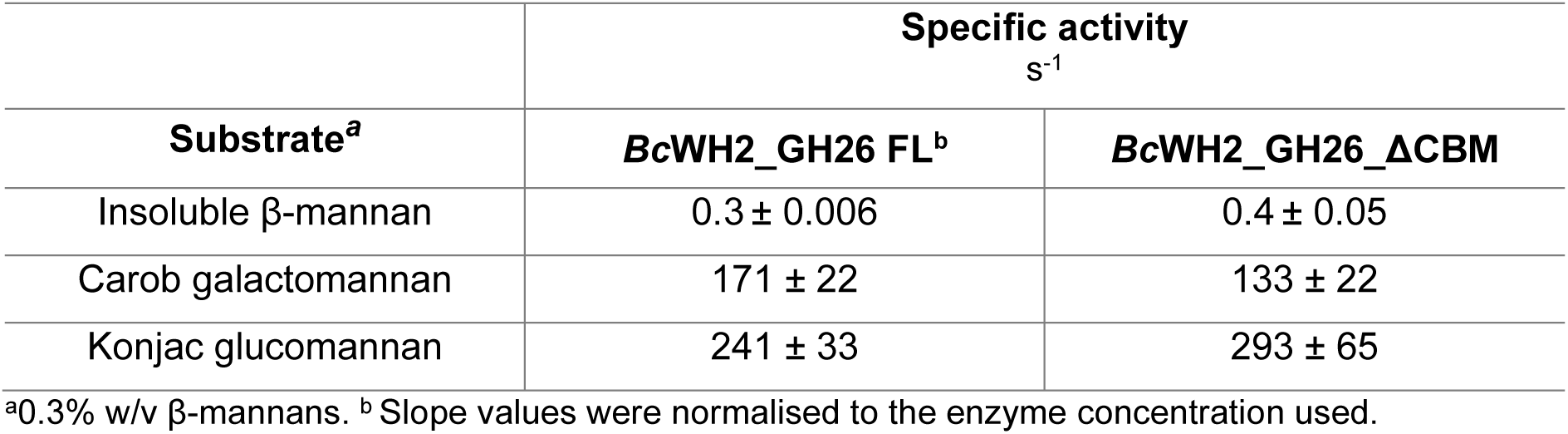
Specific activity of two *Bc*WH2_GH26 enzyme constructs on soluble and insoluble mannans.

| Substrate<sup>a</sup> | Specific activity<br>s<sup>-1</sup> |  |
| --- | --- | --- |
|  | *BcWH2_GH26 FL*<sup>b</sup> | *BcWH2_GH26_ΔCBM* |
| Insoluble β-mannan | 0.3 ± 0.006 | 0.4 ± 0.05 |
| Carob galactomannan | 171 ± 22 | 133 ± 22 |
| Konjac glucomannan | 241 ± 33 | 293 ± 65 |

<sup>a</sup>0.3% w/v β-mannans. <sup>b</sup> Slope values were normalised to the enzyme concentration used.

### Proposed ligand binding model for CBMXX

As crystallisation trials to obtain a ligand complex were unsuccessful, we used *in silico* tools to generate a model of *Bc*WH2_GH26 CBMXX bound to mannotetraose, Figure 9A. The location of the CBM binding site and the positioning of the residues in relation to the ligand are consistent with the data obtained from analysis of the CBM point mutants. In common with most CBM binding sites, tryptophan residues (W257 and W301 in *Bc*WH2_GH26 CBMXX) play a key role in ligand recognition by forming CH–π stacking interactions with the mannose rings of Man1 and Man3 (labelled from the reducing to non-reducing end), Figure 9B. The spatial arrangement of the OH-6 group of Man1 relative to W257 suggests that the addition of a galactosyl side chain at this position would introduce a steric clash, thereby impeding binding and, consequently, providing rationale for the inability of the CBM to accommodate heavily-galactosylated ligands.

**Figure 9.**
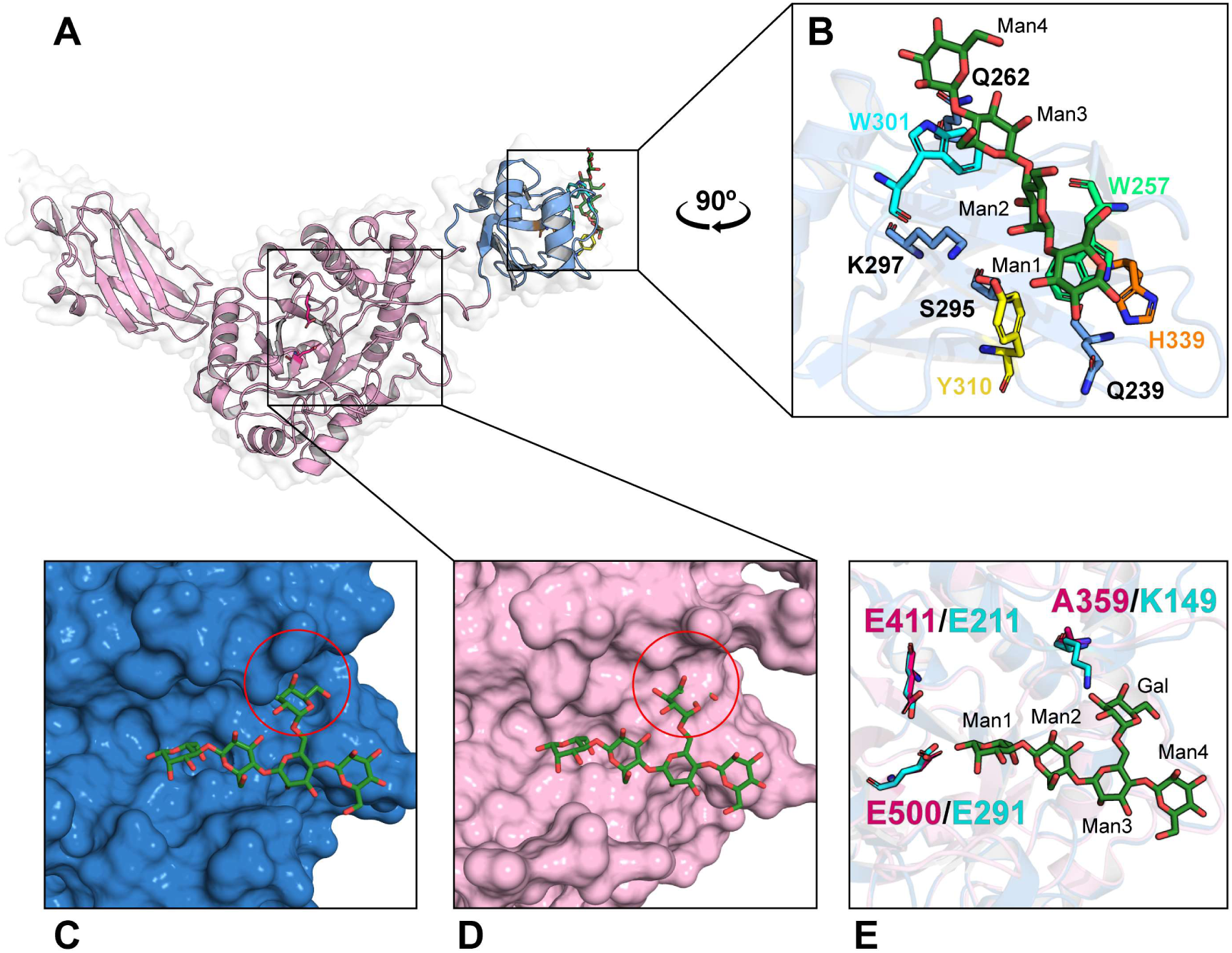
Model of *Bc*WH2_GH26 catalytic domain and CBM bound to mannooligosaccharides. A) AlphaFold2 model of a full-length enzyme with a docking prediction of its CBM (in blue) bound to linear mannotetraose (M4) created using Chai-1 Discovery software(26). B) Close-up of the CBM binding site with M4 (in green) modelled in. Binding residues highlighted. C) Crystal structure of *Bo*ManGH26B (PDB: 6HF4) active site with a decorated galactomanno-oligosaccharide (M4 with single Gal side chain at Man3; G1M4) bound. Red circle highlights structural pocket allowing accommodation of galactose side chain on Man3. D) Model of a *Bc*WH2_GH26 active site with a G1M4 from PDB 6HF4 overlaid into active site. Red circle indicates the Tyr201 residue causing steric clash with the galactose side chain. E) Superimposition of *Bc*WH2_GH26 and *Bo*ManGH26B active sites with G1M4 bound. *Bo*ManGH26B catalytic residues (E211 and E291) are shown in cyan. Also shown is Lys149 (cyan) in *Bo*GH26 that forms a hydrogen bond with the galactose side chain. Highlighted in pink are putative catalytic residues (E411 and E500) of *Bc*WH2_GH26 and alanine residue (A359) in the analogous position to *Bo* Lys149, preventing hydrogen bond formation with the galactose side chain of Man3.

Furthermore, the positioning of Y310 supports its involvement in hydrogen bonding with the sugar, most likely via interaction between its phenolic hydroxyl group and OH-2 of Man2. Such structural arrangement could impact ligand specificity at this subsite, *i.e.,* an alternative C2 hydroxyl configuration, such as equatorial OH-2 of glucose, could cause a steric clash at this subsite and thus preclude binding.

Other residues *Bc*WH2_CBM binding cleft capable of hydrogen bond formation, which could influence ligand specificity include Q262, Q293, S295, K297, and H339 (Figure 9B), the last of which has been experimentally shown to be involved in ligand binding, Figure 8, most likely via interaction with OH-1 of Man1.

Besides the two tryptophans, Q293 is the amongst the most conserved residues within the binding cleft, suggesting it might play a key role in determining ligand specificity, Figure 8A. It most likely interacts with Man1 and its location at the periphery of the binding cleft provides structural rationale for DP4 being the minimum chain length required for binding, despite the apparent suitability of mannotriose to span the two tryptophan residues (W257 and W301), Figure 9B.

To investigate the relative positioning of the CBM binding site in the context of the full length enzyme the model of the *Bc*WH2_GH26 catalytic domain was overlaid with the crystal structure of *B. ovatus Bo*Man26B (PDB: 6HF4) consisting of a singular domain (52% amino acid sequence identity shared with a catalytic domain of *Bc*WH2_GH26), bound to a decorated galactomanno-oligosaccharide (M4 with single Gal side chain at Man3; G1M4), Figure 9A. This revealed that the CBM’s binding site appears to face in the opposite direction and orientation to the catalytic domain’s binding cleft, suggesting that the CBM does not act as a direct extension of the enzyme’s active site *i.e.* appears unlikely to bind to the same mannan chain as the enzyme.

### Accommodation of decorated mannans by BcWH2-GH26 catalytic domain

Overlay of the active sites of the two GH26 enzymes also revealed some interesting differences in their ability to accommodate decorated mannans, Figure 9C-E. Although the catalytic residues (E411 and E500 for *Bc*WH2_GH26) are conserved as expected within the same family, the two enzymes differ in their predicted −4 subsites. The tyrosine residue (Y201) in *Bc*WH2_GH26 clashes with the Gal decoration on the M3 of the overlaid G1M4, as indicated by a red circle in Figure 9D. Steric clashes with potential O6 Gal decorations might also occur at the Man1 due to the presence of Y524. Based on the overlay, the galactose side chains could therefore be accommodated at Man2 and Man4, but they cannot be localised on two neighbouring mannose residues.

Furthermore, in the case of *Bo*Man26B, K149 forms hydrogen bonds with a galactosyl-side chain of Man3(22), whereas, in *Bc*WH2_GH26, this residue is substituted with alanine suggesting no direct interaction between the *Bc*WH2_GH26 and galactose at this site.

Overall, these structural features preventing binding of galactosyl-side chains at multiple subsites in *Bc*WH2_GH26, confer the enzyme unable to act upon highly decorated regions of galactomannans. This is thus consistent with the previously observed inability of its CBM to bind highly galactosylated ligands.

However, all of this analysis is based on modelled sugars in the CBM and sugar from an overlay with the catalytic domain and thus caution must be taken with the conclusions drawn.

## Discussion

In this study, we functionally characterised two founding members of a novel CBM family – CBMXX. CBMXXs tested in this study are type B CBMs, able to bind soluble and insoluble ligands, most likely by interacting with the non-crystalline regions of the latter(27). Interestingly, both CBMXXs exhibited very narrow range of specificity, binding solely to β-mannan, and failing to interact with other β-linked polysaccharides tested, Table 1. This stands in contrast with many other characterised β-mannan binding CBMs, which display higher degree of ligand promiscuity. For instance, members of family CBM29, are able to bind β-mannan, but also β-linked glucans and cellulose, due to very few hydrogen bonds being formed between their binding site residues and the C2 hydroxyl of the ligand, Figure S11 (28). In contrast, *Bc*WH2_CBM binding site contains multiple residues capable of forming hydrogen bonds (H339, Y310, Q262, Q293, K297, S295), and their importance in ligand binding has been confirmed via affinity studies of targeted mutants (Y310 and H339), as per Figure 8. We speculate, based on the Chai1 binding model, Figure 9, that the positioning of Y310 could potentially limit the ligand specificity of the CBM, preventing binding of glucose at this subsite, with β1,4-linked mannosyl-saccharides as the only conformationally compatible ligands. This is further supported by ITC data, which show that *Bc*WH2_CBM is unable to bind G5 despite binding M5, highlighting the likely involvement of C2 hydroxyl configuration in ligand specificity. Nevertheless, *Bc*WH2_CBM and *Bu*_CBM were able to bind glucomannan, a polymer consisting of both glucose and mannose residues. These observations indicate that the CBMs either exhibit specificity for backbone regions composed exclusively of mannose residues, or alternatively, they demonstrate a capacity to accommodate glucose residues at certain subsites, provided that mannose occupies the critical positions essential for binding. This represents a distinct recognition mechanism compared to the previously characterized glucomannan-binding CBM16 and CBM29 families, which can accommodate either hexose residue within the key binding sites, as demonstrated by their capacity to bind both cello- and manno-oligosaccharides, Figure S11 (29, 30).

Moreover, both CBMXX modules tested failed to bind 6³,6⁴-di-galactosyl-mannopentaose (G2M5), despite binding to CGM, Table 1. This observation further supports the notion that their capacity to accommodate main chain substitutions is restricted only to specific subsites within the binding cleft. For instance, based on the Chai1 model, we suspect that the positioning of W257 might preclude accommodation of galactosyl-side chain at Man1. This is not unusual and has been previously observed within the binding site of CBM27, in which the galactosyl-substitutions can only be tolerated in subsites 1, 2, 3 and 5, consequently, explaining this CBMs low affinity for G2M5, Figure S11 (13). CBMXX displaying a similar pattern of discrimination of side chains to CBM27 agrees with the observed lower affinity for the highly galactosyl-decorated GGM compared to CGM, as shown by native affinity gels, Figure S2. This specificity for relatively unsubstituted galactomannan regions is mirrored in the BcWH2_GH26 catalytic domain. Comparison between predicted model of *Bc*WH2_GH26 and crystal structure of *Bo*GH26B, suggests that the *Bc*WH2_GH26 active site might be unable to accommodate galactosyl-side chains, due to a steric hindering caused by Tyr210, Figure 9. Combining the experimental data with the computational structural predictions, we speculate that although *Bc*WH2_GH26 and its CBM are able to target galactomannans, they can only access less decorated regions of these storage polysaccharides. A preference for low-substituted galactomannan substrates has previously been described in other mannanases, such as *Bacteroides ovatus* GH26A enzyme (*Bo*Man26A) which is thought to act in synergy with a GH36 α-galactosidase from the same PUL to degrade galactosylated substrates(2). In contrast to *B. ovatus* β-mannan PUL, *Bc*PUL32 does not contain a GH36, however, a non-PUL GH36 gene was found to be highly upregulated during *B. cellulosilyticus* growth on galactomannan(16).

One of the most interesting aspects of the two CBMs tested in this study is their unusual positioning within the enzyme. Although uncommon, the existence of internal CBMs has been previously reported in several GH families, such as GH10 xylanases(31), GH13 amylases(32), and GH148 glucanases (33). Most recently, a structural homolog of *Bc*WH2_GH26 has been identified, however, no functional characterisation of its putative internal CBM had been undertaken(17). Despite this, the rationale for evolving such molecular architecture over the more classical terminal CBM arrangement remains poorly understood.

To determine the CBMs biological role, enzymatic activity of the full-length *Bc*WH2_GH26 and *Bc*WH2_GH26_ΔCBM was compared. Interestingly, no significant differences were observed between the two constructs when comparing their activity on either soluble or insoluble substrates (Table 3). The lack of difference in activity against insoluble mannan indicates that *Bc*WH2_CBM might not play the canonical role of CBMs, which enhance catalytic performance by targeting enzymes to the hard-to-access polysaccharides embedded within the plant cell walls. Nonetheless, the ability of *Bc*WH2_CBM to bind insoluble mannan suggests its potential involvement in targeting of insoluble substrates in an alternative way. Considering the human gut environment and its short transit time compared to, for example, ruminants, the CBM might be an adaptation for efficient degradation of insoluble fibres derived from our diet. The lack of an effect on the activity of its corresponding enzyme could be attributed to the nature of the assay itself, which fails to mimic the true gut environment in which the enzyme functions.

Furthermore, the obtained results do not provide explanation for the unique positioning of the CBM within the *Bc*WH2_GH26 catalytic domain. To fully understand the function of this CBM and rationale for developing such a unique molecular architecture, broader context of an outer membrane needs to be considered. Based on the composition of *Bc* PUL32, *Bc*WH2_GH26, alongside either one or both SGBPs, most likely forms part of a β-mannan utilisome(*34*). The location of the CBM, as well as the presence of the N-terminal β-sheet domain might therefore both be structural adaptations helping position the enzyme within utilisome complex. Cryo EM structural studies of the intact mannan utilisome could provide invaluable insight into the spatial positioning of the CBM relative to the other utilisome components and their binding sites and elucidate the mechanistic basis of their potential interplay.

The CBM internal positioning as a utilisome adaptation is further evidenced when comparing other enzymes possessing similar domain architecture. Both *B. thetaiotaomicron* SusG and *Pseudacanthotermes militaris* GH10 contain internal CBMs and are located within PULs containing SGBPs(31, 35), strongly suggesting they form complexes analogous to *B. cellulosilyticus* β-mannan utilisome. Since internal CBMs are found within various enzyme classes, they are most likely not a polysaccharide-specific adaptation. Nonetheless, they likely function to improve the catalytic efficiency of the utilisation system, providing bacterium with competitive advantage when growing on a specific polysaccharide.

To confirm whether all CBMXX modules function in a context of a utilisome, we performed genomic analysis of the family. We found that 61 out of 82 proteins associated with these CBMs are GH26s of a structure homologous to *Bc*WH2_GH26, *i.e.,* that contain both the N-terminal extension and an internal CBM. Majority of them are located within PULs and therefore, most likely function within a utilisome.

This notion is further reinforced by the instances in which the CBMXX is not associated with a GH26, and instead, serves as the glycan binding domain of putative SGBPs, such as B5F25_13220 or Bacsa_1712, Figure 6B. These SGBPs are located within PULs also containing GH26 genes. The GH26s (*e.g.,* B5F25_13225 or Bacsa_1713) are structural homologues of *Bc*WH2_GH26, but do not contain the internal CBM. Provided that these proteins are all components of the same outer membrane utilisomes this could indicate that the role of CBMXX depends on its specific location in the utilisome rather than in the GH26 per se. It should be noted, however, that this hypothesis does not extend to all members of the CBMXX family, as the present discussion focuses specifically on the domains associated with GH26s and SGBPs localised in PUL. A subset of CBMXX is found associated with proteins predicted to be periplasmic rather than localised to the outer membrane, such as CE7s, in which case the CBM likely plays a different role.

Overall, the results of this study provide initial insight into novel β-mannan active enzymes and their function within complex degradation systems. These insights contribute to our understanding of glycan breakdown by gut microbes, essential for the development of novel, diet-based, health-promoting interventions.

Furthermore, as β-mannans are recognised contributors to household stains (36), the unique architecture of the enzyme might be useful when exploited in industrial applications such as fabric and home-care.

