## Supplementary Information for "Unusual molecular architecture of a human gut microbiota β-mannanase reveals a new CBM family"

**Table S1. Bacteroidota media solutions and supplements.**

| Supplement name | Component name | Final conc. |
|---|---|---|
| **10x Bacteroidetes salts** | $KH_2PO_4$ | 136 g/l |
| | NaCl | 8.75 g/l |
| | $(NH_4)_2SO_4$ | 11.25 g/l |
| | pH adjusted to 7.2 and filter sterilised with sterile 0.2 μm filter. | |
| **Balch's vitamins** | *p*-Aminobenzoic acid | 5 mg/l |
| | Folic acid | 2 mg/l |
| | Biotin | 2 mg/l |
| | Nicotinic acid | 5 mg/l |
| | Calcium pantothenate | 5 mg/l |
| | Riboflavin | 5 mg/l |
| | Thiamine HCl | 5 mg/l |
| | Pyridoxine HCl | 10 mg/l |
| | Cyanocobalamin | 0.1 mg/l |
| | Thioctic acid | 5 mg/l |
| | pH adjusted to 7.0 and filter sterilised with sterile 0.2 μm filter. | |
| **Trace mineral supplement** | EDTA | 0.5 g/l |
| | $MgSO_4*7H_2O$ | 3 g/l |
| | $MnSO_4*H_2O$ | 0.5 g/l |
| | NaCl | 1 g/l |
| | $FeSO_4*7H_2O$ | 0.1 g/l |
| | $CaCl_2$ | 0.1 g/l |
| | $ZnSO_4*7H_2O$ | 0.1 g/l |
| | $CuSO_4*5H_2O$ | 0.01 g/l |
| | $H_3BO_3$ | 0.01 g/l |
| | $Na_2MoO_4*2H_2O$ | 0.01 g/l |
| | $NiCl_2*6H_2O$ | 0.02 g/l |
| | pH adjusted to 7.0 and filter sterilised with sterile 0.2 μm filter. | |
| **Purine/pyrimidine solution** | Adenine | 200 mg/l |
| | Guanine | 200 mg/l |
| | Thymine | 200 mg/l |
| | Cytosine | 200 mg/l |
| | Uracil | 200 mg/l |
| | pH adjusted to 7.0 and filter sterilised with sterile 0.2 μm filter. | |
| **Amino acid solution** | Alanine | 250 mg/l |
| | Arginine | 250 mg/l |
| | Asparagine | 250 mg/l |
| | Aspartic acid | 250 mg/l |
| | Cysteine | 250 mg/l |
| | Glutamic acid | 250 mg/l |
| | Glutamine | 250 mg/l |
| | Glycine | 250 mg/l |
| | Histidine | 250 mg/l |
| | Isoleucine | 250 mg/l |
| | Leucine | 250 mg/l |
| | Lysine | 250 mg/l |
| | Methionine | 250 mg/l |
| | Phenylalanine | 250 mg/l |
| | Proline | 250 mg/l |
| | Serine | 250 mg/l |
| | Threonine | 250 mg/l |
| | Tryptophan | 250 mg/l |
| | Tyrosine | 250 mg/l |
| | Valine | 250 mg/l |
| | pH adjusted to 7.0 and filter sterilised with sterile 0.2 μm filter. | |

**Table S2. Bacteroidota minimal medium (MM) composition.**

| Component name[a] | Volume added |
|---|---|
| 10x Bacteroidetes salts | 10 ml |
| Balch's vitamins | 1 ml |
| Trace mineral supplement (1 mg/ml) | 1 ml |
| Purine/pyrimidine solution (1 mg/ml) | 1 ml |
| Amino acid solution (5 mg/ml) | 1 ml |
| Vitamin K3 (1 mg/ml) | 100 µl |
| $FeSO_4$ (0.4 mg/ml) | 100 µl |
| $CaCl_2$ (0.8% w/v) | 100 µl |
| $MgCl_2$ (0.1 M) | 100 µl |
| Hematin (1.9 mM)-histidine (0.2M) | 100 µl |
| Vitamin B12 (0.01 mg/ml) | 50 µl |
| L-cysteine | 100 mg |
| $dH_2O$ | 35.85 ml |

[a]pH adjusted to 7.2 and filter sterilised with sterile 0.2 µm filter.

**Table S3. Bacteroidota species used in this study.**

| No. | Species name | Strain identifier |
|---|---|---|
| **1** | *Bacteroides caccae* | ATCC 43185 |
| **2** | *Bacteroides cellulosilyticus* | WH2 |
| **3** | *Bacteroides clarus* | DSM 22519 |
| **4** | *Phocaeicola dorei* | DSM 17855 |
| **5** | *Bacteroides eggerthii* | DSM 20697 |
| **6** | *Bacteroides finegoldii* | DSM 17565 |
| **7** | *Bacteroides fluxus* | DSM 22534 |
| **8** | *Bacteroides fragilis* | NCTC 9343 |
| **9** | *Bacteroides intenstinalis* | DSM 17393 |
| **10** | *Phocaeicola massiliensis* | DSM 17679 |
| **11** | *Bacteroides oleiciplenus* | DSM 22535 |
| **12** | *Bacteroides ovatus* | ATCC 8483 |
| **13** | *Phocaeicola plebeius* | DSM 17135 |
| **14** | *Bacteroides salyersiae* | DSM 18765 |
| **15** | *Bacteroides stercoris* | ATCC 43183 |
| **16** | *Bacteroides thetaiotaomicron △TDK* | ATCC 29148 |
| **17** | *Bacteroides uniformis* | ATCC 8492 |
| **18** | *Phocaeicola vulgatus* | ATCC 8482 |
| **19** | *Bacteroides xylanisolvens* | XB1A |

**Table S4. Primers used in this study.**

| Construct | Primer Name | Sequence[a] |
|---|---|---|
| *Bc*WH2_GH26 FL | *Bc*WH2_GH26_NheI_FP | CTC GCT AGC TGC ACT GAT ACT GAT GCA CAA TAT AC |
| | *Bc*WH2_GH26_XhoI_RP | CTC CTC GAG CTT CAT GTC AGG TAA GTC CTC |
| *Bc*WH2_GH26 ΔN-term | *Bc*WH2_GH26_NheI_FP | CTC GCT AGC TTG GAT AAA TCG TTG GTA AAT GCT TC |
| *Bu*_CBM | *Bu*_CBM_NheI_FP | CTC GCT AGC GGC GAA GCT TCT TCT ACC |
| | *Bu*_CBM_XhoI_RP | CTC CTC GAG TGT ACC GTT ATT TTC AAG ATA AAC AG |

[a]Restriction sites are underlined.

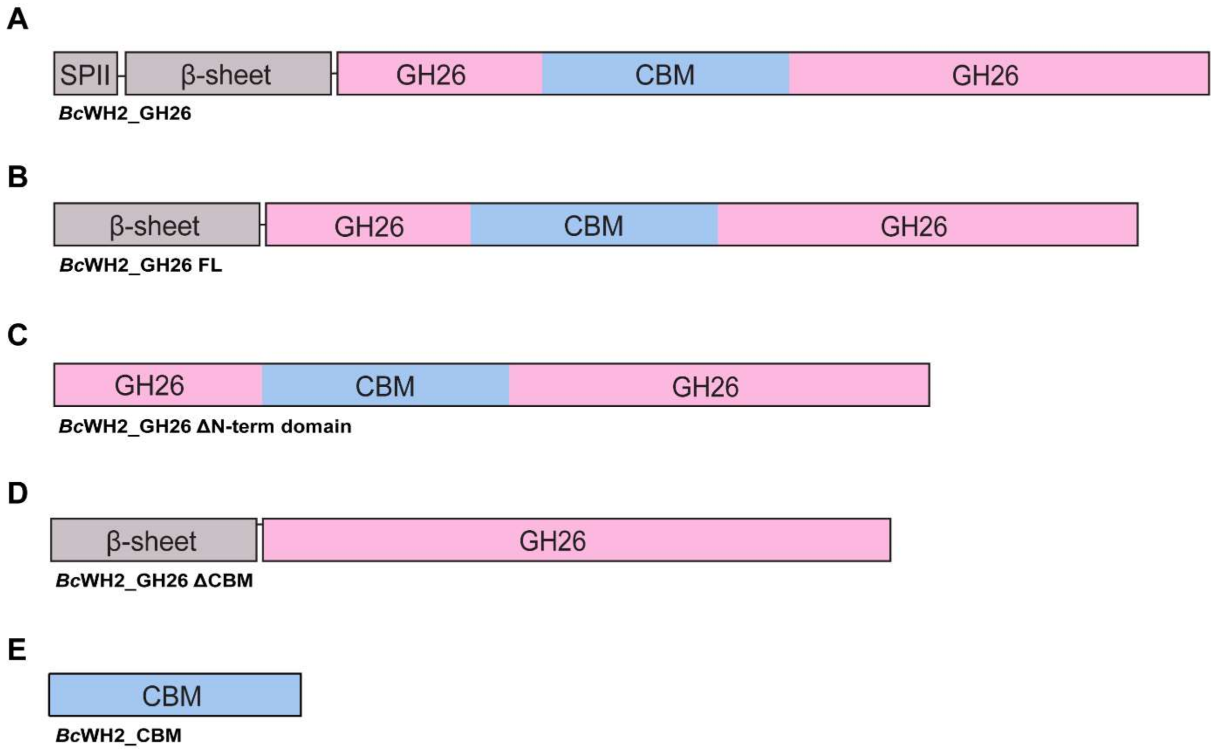


**Figure S1. Schematic of *Bc*WH2_GH26 constructs used in the study.** A) Wild-type, full-length protein. B) Full-length mature construct lacking signal peptide. C) *Bc*WH2_ΔN-term domain. D) *Bc*WH2_ΔCBM. E) *Bc*WH2_CBM.

**Table S5. Sequences of constructs synthesised by Twist Biosciences and used in this study.**

| Construct | | Sequence[a] |
|---|---|---|
| *Bc*WH2_GH26 ΔCBM[b] | DNA | TGCACTGATACTGATGCACAATATACCATTCCCGAAGTGGAGGCACCTGTACTGGTGTCCACCA<br>CTCCGGAGGCGAATGCTGCCAAGGTGAAGAGAGGCGAAATCACCATTGAGGTGAAGTATGAC<br>AAGAACGTGTTCTTTGCTACGGAAGACTTGAAGCAAATTTCGTTTACGGGTGGCACGCTGATTA<br>GTGCTGATGTATATGGGTCCAGTAATGTACTGACACTGACTGTAGAAGTACCCACACGCGAAAC<br>TGTGTGTTCTCTCTCTATTCCCGAAGGTGTAGTGCTGGGGCCGAATAAGACGCCTGCTCCGGC<br>TGTATCTCTTCAGTTTACTACCGTTGCTTTGGATAAATCGTTGGTAAATGCTTCTGCTACTCCTGC<br>TGCACAGAAAGTTTATGCATATTTGTTAGAAAACTTTGAGAGCAAGACTCTTTCGGCTATGATGG<br>CCAATGTGAATTGGAATACGGAAAAGTCGGAACAGGTATATCAGTGGACAGGTAAGTATCCTGC<br>TATCAATTGTTTTGACTATGTGCATCTGTATGCTTCTGGTGCAAACTGGATTAACTATAGTGATATT<br>ACTCCTGTGAAAGATTGGTGGAATGCCGGTGGTTTAGTGTCTGCTATGTGGCATTGGAATGTGC<br>CGACAAAGGCAACTAAATATGCATTCTATAGGGAAGATACGGATTTTGATGCAGACAATGCTTTG<br>ACTGAGGGAACTTGGGAGAATAAAGTCTTTACTGAAGATTTGGCTAAAGTTGCGGCTAATTTAAA<br>ATTGTTGCAAGATGAAGGCATTCCTGTAATTTGGCGTCCATTCCATGAGGCTGCCGGTGGCTG<br>GTTCTGGTGGGGTAAGAATGCCACAAGCTTCAAGAATATGTGGATTGCCATGTTCAACTACTTTA<br>AGGCAGAAGGGTTGAATAACCTGATCTGGGTATGGACTACTGAAACCGGTGATGACGATTGGTA<br>TCCGGGTGATGCTTATGTGGATATTGTGGGACGTGATATTTATACGAAAGATGCAAGCACTTGTG<br>CTTCAGATTATAGTTCTATAGTAGTAGCTTATGGCAATAAGATGGTTGCATTAAGCGAGTGTGGCA<br>CGGTAGGCAAGATTTCCGAGCAATGGGCAGCCGGTGCACGCTGGTCTTGGTTTATGCCTTGGT<br>ATGACGCGGAAGACGCGGAAACGCCACATGCTGATCAGGCTTGGTGGAAAGATGCTATGGAA<br>CAAAATTTCGTGATATCCCGTGAGGACTTACCTGACATGAAGTAG |
| | Protein | MGSSHHHHHHSSGLVPRGSHMCTDTDAQYTIPEVEAPVLVSTTPEANAAKVKRGEITIEVKYDKNV<br>FFATEDLKQISFTGGTLISADVYGSSNVLTLTVEVPTRETVCSLSIPEGVVLGPNKTPAPAVSLQFTTV<br>ALDKSLVNASATPAAQKVYAYLLENFESKTLSAMMANVNWNTEKSEQVYQWTGKYPAINCFDYVH<br>LYASGANWINYSDITPVKDWWNAGGLVSAMWHWNVPTKATKYAFYREDTDFDADNALTEGTWEN<br>KVFTEDLAKVAANLKLLQDEGIPVIWRPFHEAAGGWFWWGKNATSFKNMWIAMFNYFKAEGLNNL<br>IWVWTTETGDDDWYPGDAYVDIVGRDIYTKDASTCASDYSSIVVAYGNKMVALSECGTVGKISEQ<br>WAAGARWSWFMPWYDAEDAETPHADQAWWKDAMEQNFVISREDLPDMK |
| *Bc*WH2_CBM WT[b] | DNA | CCTGATGTTTTCTCTGAGGGGCTATGGACGGGCGAACAGGCTATGCCCGGTGATTGGAGTGGA<br>AATGTTCAGTTGACGGATGATGCAGCAAAAGTTGCTTTTGCTGAGGCACAAGTGGGGAATAAA<br>GTAAGAGTAACTGTAAAAGATATAGCTGCGGGTGCACAAGGCTCTTTTAAAAATAGTAGCTGGCT<br>TGAAATCGCTCCAGGTATGGATTATTTTGATATTACAGGTGACTTTGAATTGGAGATTACCGAAG<br>CTGTTCTGACATCTCTGAAAGATGGTGGTTTAATTATCGGTGGTCACGACTATACGGTTACTGGC<br>GTTTATTTGGAAGGTGGTTCGGCAAGTGAT |
| | Protein | MGSSHHHHHHSSGLVPRGSHMPDVFSEGLWTGEQAMPGDWSGNVQLTDDAAKVAFAEAQVGN<br>KVRVTVKDIAAGAQGSFKNSSWLEIAPGMDYFDITGDFELEITEAVLTSLKDGGLIIGGHDYTVTGVY<br>LEGGSASDLEHHHHHH |
| *Bc*WH2_CBM W257A[b] | DNA | CCTGATGTTTTCTCTGAGGGGCTATGGACGGGCGAACAGGCTATGCCCGGTGATGCCAGTGG<br>AAATGTTCAGTTGACGGATGATGCAGCAAAAGTTGCTTTTGCTGAGGCACAAGTGGGGAATAAA<br>GTAAGAGTAACTGTAAAAGATATAGCTGCGGGTGCACAAGGCTCTTTTAAAAATAGTAGCTGGCT<br>TGAAATCGCTCCAGGTATGGATTATTTTGATATTACAGGTGACTTTGAATTGGAGATTACCGAAG<br>CTGTTCTGACATCTCTGAAAGATGGTGGTTTAATTATCGGTGGTCACGACTATACGGTTACTGGC<br>GTTTATTTGGAAGGTGGTTCGGCAAGTGAT |
| | Protein | MGSSHHHHHHSSGLVPRGSHMPDVFSEGLWTGEQAMPGD**A**SGNVQLTDDAAKVAFAEAQVGNK<br>VRVTVKDIAAGAQGSFKNSSWLEIAPGMDYFDITGDFELEITEAVLTSLKDGGLIIGGHDYTVTGVYL<br>EGGSASDLEHHHHHH |
| *Bc*WH2_CBM W301A[b] | DNA | CCTGATGTTTTCTCTGAGGGGCTATGGACGGGCGAACAGGCTATGCCCGGTGATTGGAGTGGA<br>AATGTTCAGTTGACGGATGATGCAGCAAAAGTTGCTTTTGCTGAGGCACAAGTGGGGAATAAA<br>GTAAGAGTAACTGTAAAAGATATAGCTGCGGGTGCACAAGGCTCTTTTAAAAATAGTAGCGCCCT<br>TGAAATCGCTCCAGGTATGGATTATTTTGATATTACAGGTGACTTTGAATTGGAGATTACCGAAG<br>CTGTTCTGACATCTCTGAAAGATGGTGGTTTAATTATCGGTGGTCACGACTATACGGTTACTGGC<br>GTTTATTTGGAAGGTGGTTCGGCAAGTGAT |
| | Protein | MGSSHHHHHHSSGLVPRGSHMPDVFSEGLWTGEQAMPGDWSGNVQLTDDAAKVAFAEAQVGN<br>KVRVTVKDIAAGAQGSFKNSS**A**LEIAPGMDYFDITGDFELEITEAVLTSLKDGGLIIGGHDYTVTGVY<br>LEGGSASDLEHHHHHH |
| *Bc*WH2_CBM Y310A[b] | DNA | CCTGATGTTTTCTCTGAGGGGCTATGGACGGGCGAACAGGCTATGCCCGGTGATTGGAGTGGA<br>AATGTTCAGTTGACGGATGATGCAGCAAAAGTTGCTTTTGCTGAGGCACAAGTGGGGAATAAA<br>GTAAGAGTAACTGTAAAAGATATAGCTGCGGGTGCACAAGGCTCTTTTAAAAATAGTAGCTGGCT<br>TGAAATCGCTCCAGGTATGGATGCCTTTGATATTACAGGTGACTTTGAATTGGAGATTACCGAAG<br>CTGTTCTGACATCTCTGAAAGATGGTGGTTTAATTATCGGTGGTCACGACTATACGGTTACTGGC<br>GTTTATTTGGAAGGTGGTTCGGCAAGTGAT |
| | Protein | MGSSHHHHHHSSGLVPRGSHMPDVFSEGLWTGEQAMPGDWSGNVQLTDDAAKVAFAEAQVGN<br>KVRVTVKDIAAGAQGSFKNSSWLEIAPGMD**A**FDITGDFELEITEAVLTSLKDGGLIIGGHDYTVTGVY<br>LEGGSASDLEHHHHHH |
| *Bc*WH2_CBM H339A[b] | DNA | CCTGATGTTTTCTCTGAGGGGCTATGGACGGGCGAACAGGCTATGCCCGGTGATTGGAGTGGA<br>AATGTTCAGTTGACGGATGATGCAGCAAAAGTTGCTTTTGCTGAGGCACAAGTGGGGAATAAA<br>GTAAGAGTAACTGTAAAAGATATAGCTGCGGGTGCACAAGGCTCTTTTAAAAATAGTAGCTGGCT<br>TGAAATCGCTCCAGGTATGGATTATTTTGATATTACAGGTGACTTTGAATTGGAGATTACCGAAG<br>CTGTTCTGACATCTCTGAAAGATGGTGGTTTAATTATCGGTGGTGCCGACTATACGGTTACTGG<br>CGTTTATTTGGAAGGTGGTTCGGCAAGTGAT |
| | Protein | MGSSHHHHHHSSGLVPRGSHMPDVFSEGLWTGEQAMPGDWSGNVQLTDDAAKVAFAEAQVGN<br>KVRVTVKDIAAGAQGSFKNSSWLEIAPGMDYFDITGDFELEITEAVLTSLKDGGLIIGG**A**DYTVTGVY<br>LEGGSASDLEHHHHHH |

[a] Mutated residues have been highlighted in red.
[b] Construct cloned into a pET-28a(+) vector.

**Table S6. Composition of mannan activated *Bc*WH2 PUL32 (BcellWH2_02032-02020).**

| Locus tag | Protein name/CAZy family[a] | Predicted function[b] | Signal peptide[c] |
|---|---|---|---|
| BcellWH2_02020 | Est | GDSL-like Lipase/Acylhydrolase | Type I |
| BcellWH2_02021 | EPI | Cellobiose 2-epimerase | None |
| BcellWH2_02022 | MFS | Inner membrane symporter YicJ | None |
| BcellWH2_02023 | GH130_1[d] | 4-O-beta-D-mannosyl-D-glucose phosphorylase | None |
| BcellWH2_02024 | GH26 | Mannan endo-1,4-beta-mannosidase | Type II |
| BcellWH2_02025 | GH26 | Mannan endo-1,4-beta-mannanase | Type II |
| BcellWH2_02026 | Unk | Hypothetical protein | Type II |
| BcellWH2_02027 | Unk | IPT/TIG domain protein | Type II |
| BcellWH2_02028 | SusD | SusD family protein | Type II |
| BcellWH2_02029 | SusC | TonB dependent receptor | Type II |
| BcellWH2_02030 | GH5_2[d] | Endoglucanase | Type II |
| BcellWH2_02031 | GH5_7[d] | Beta-mannosidase | Type II |
| BcellWH2_02032 | CE7 | Acetyl esterase Axe7A precursor | Type I |
| BcellWH2_02033 | Unk | HTH-type transcriptional activator Btr | None |
| BcellWH2_02034 | GH3 | Beta-glucosidase | Type II |

[a] Imported from PULDB(1).
[b] From PULDB annotation, or determined based on the known activities within a given CAZy family(2).
[c] Predicted via SignalP 6.0(3).
[d] Underscored number denotes subfamily.

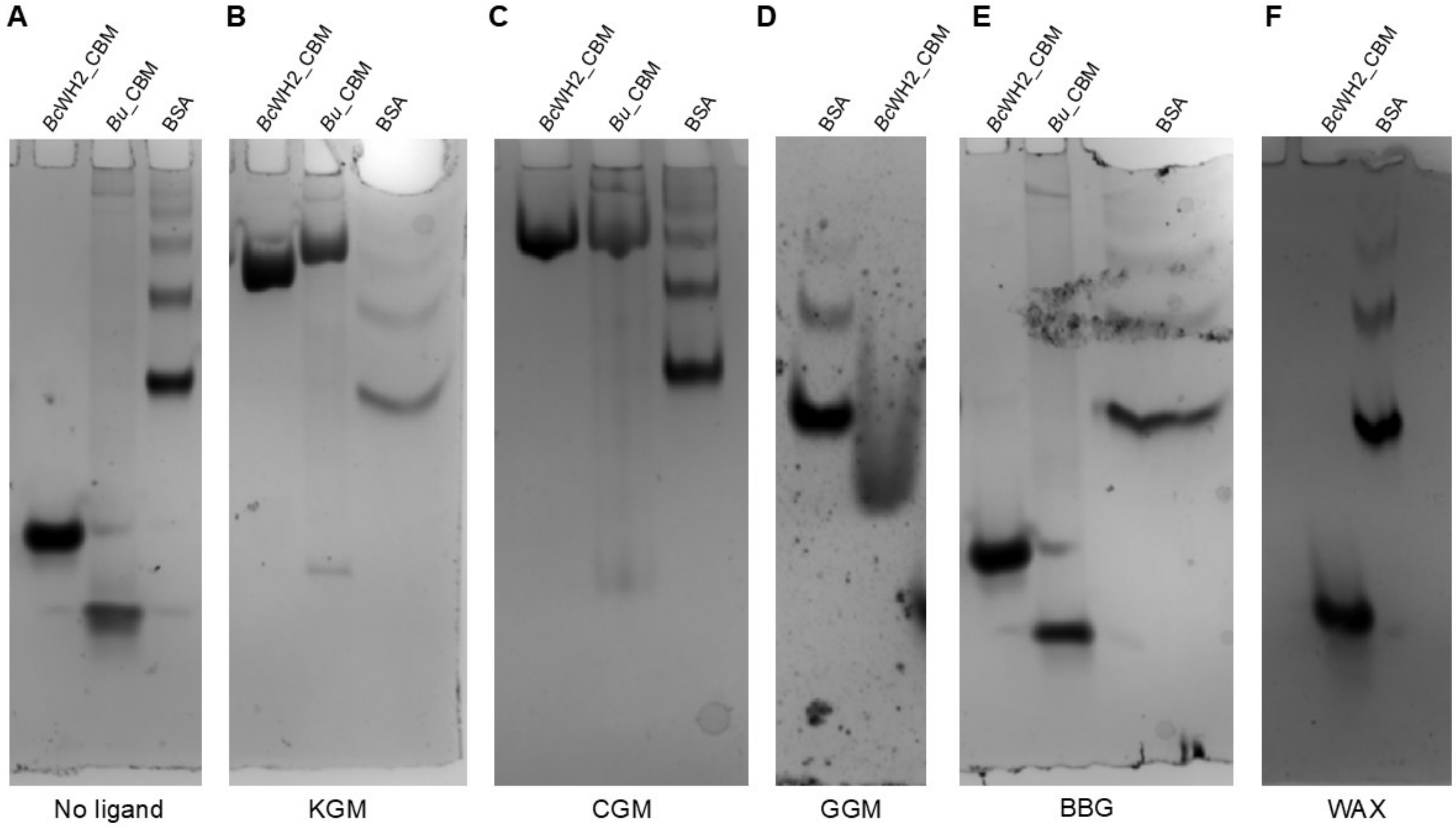


**Figure S2. Native affinity gel electrophoresis of BcWH2_CBM and Bu_CBM against various soluble polysaccharides**. A) konjac glucomannan (KGM), B) carob galactomannan (CGM), C) guar gum galactomannan (GGM), D) mixed-linkage barley β-glucan (BBG), and E) wheat arabinoxylan (WAX). Native PAGE gels were prepared using 1 mg/ml final concentration of each ligand. No ligand gel did not contain any polysaccharide and was used as a negative control. BSA was run alongside CBMs as a non-polysaccharide binding negative control.

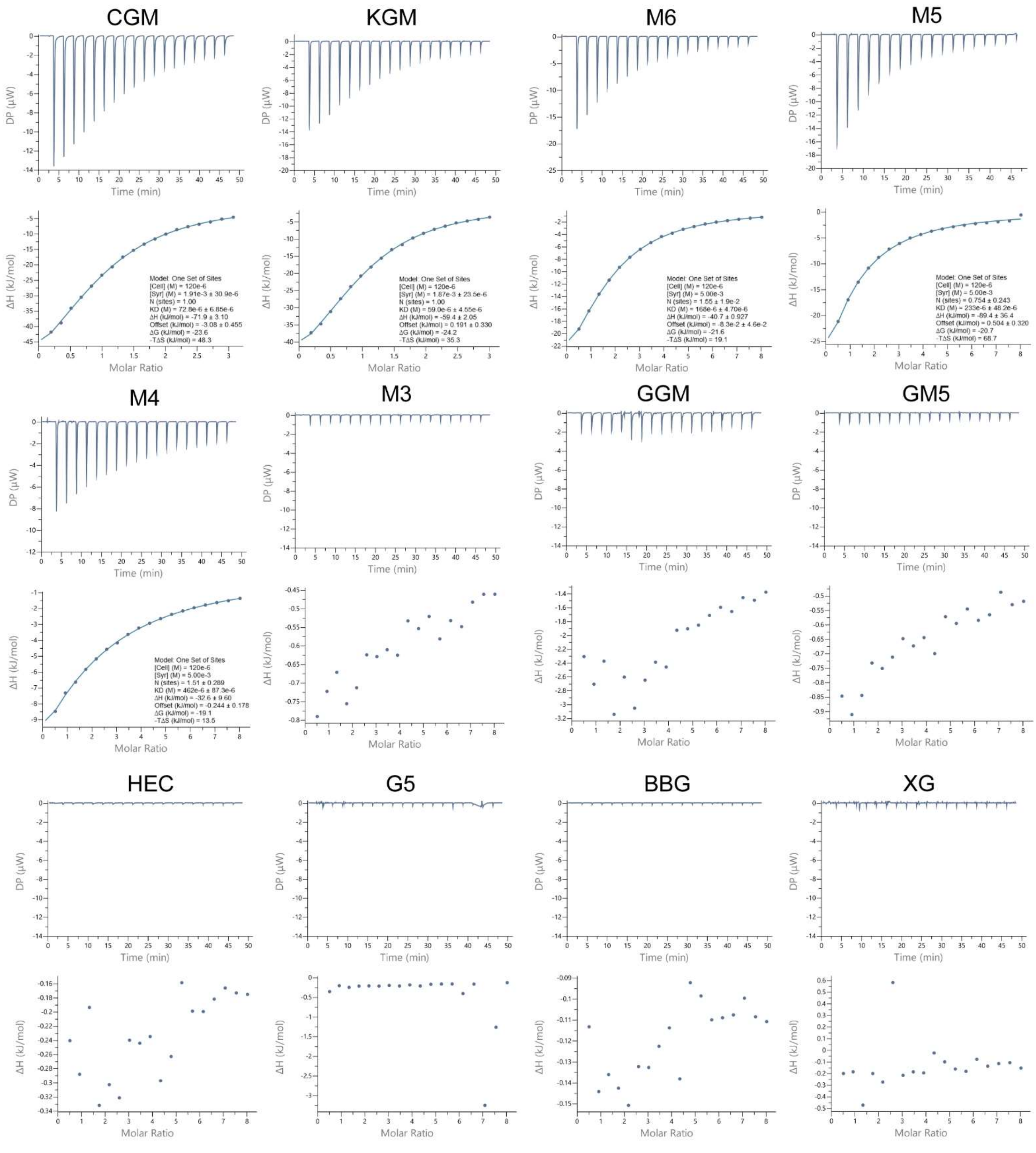


**Figure S3**. **Interactions of *Bc*WH2_CBM with various soluble ligands measured via isothermal titration calorimetry**. Titrations were carried out in 20 mM Tris buffer (pH8.0) with 150 mM NaCl added. For each ligand both raw data (top panel) and integrated data (bottom panel). The ligands used from the top left were: carob galactomannan (CGM), konjac glucomannan (KGM), mannohexaose (M6), mannopentaose (M5), mannotetraose (M4), mannotriose (M3). Then from the bottom left: guar gum galactomannan (GGM), di-galactosyl-mannopentaose (GM5), hydroxyethyl cellulose (HEC), cellopentaose (G5), mixed-linkage barley β-glucan (BBG), and xyloglucan (XG).

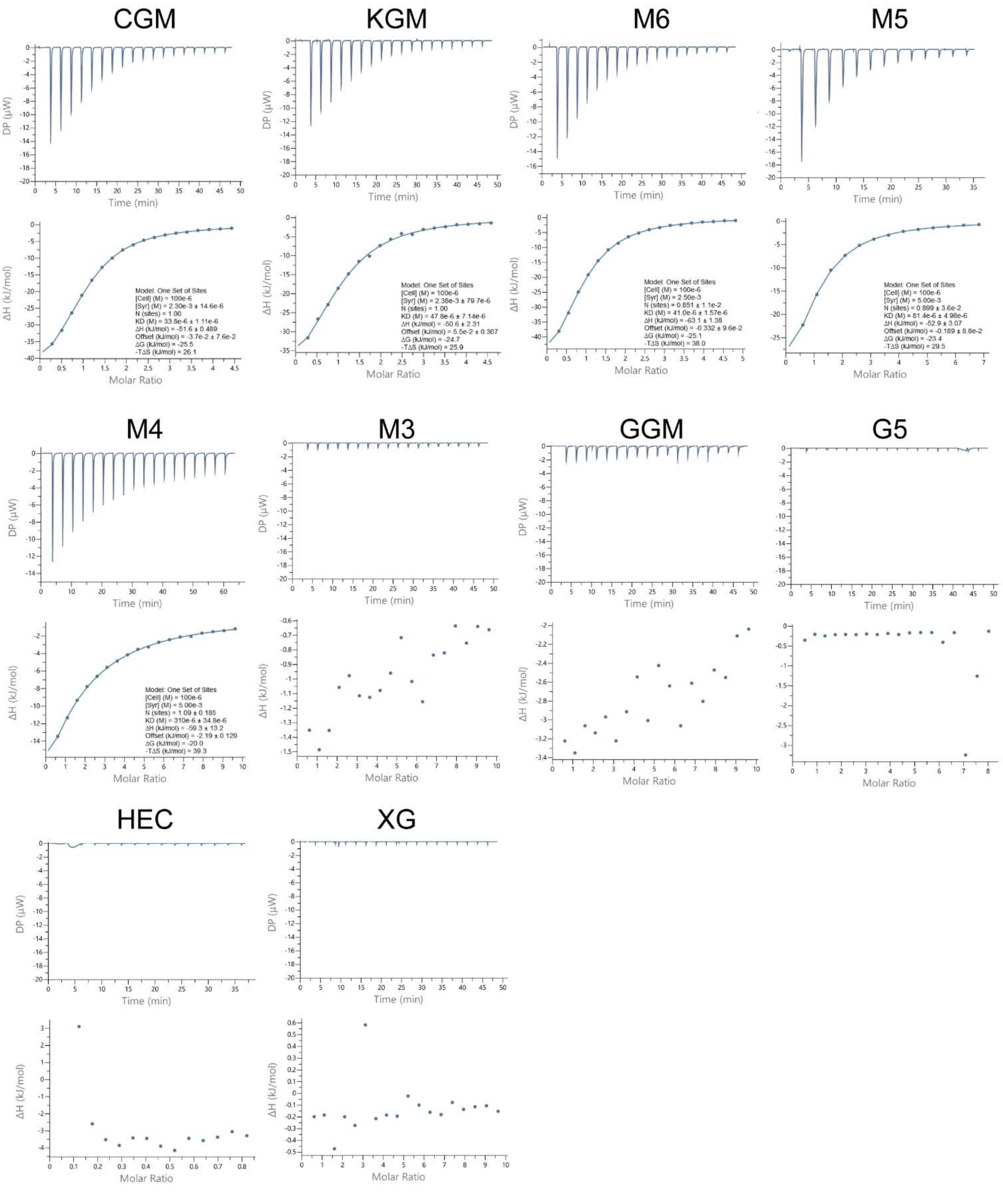


**Figure S4. Interactions of *Bu*_CBM with various soluble ligands measured via isothermal titration calorimetry**. Titrations were carried out in 20 mM Tris buffer (pH8.0) with 150 mM NaCl added. For each ligand both raw data (top panel) and integrated data (bottom panel). The ligands used from the top left were: carob galactomannan (CGM), konjac glucomannan (KGM), mannohexaose (M6), mannopentaose (M5), mannotetraose (M4). Then from the bottom left: mannotriose (M3), guar gum galactomannan (GGM), cellopentaose (G5), hydroxyethyl cellulose (HEC), and xyloglucan (XG).

**Table S7. ITC parameters determined for *Bc*WH2_CBM and *Bu*_CBM[a].**

| Construct | Ligand | $K_D$ (μM) ± SD | N ± SD | ΔG (kJ/mol) ± SD | ΔH (kJ/mol) ± SD | -TΔS (kJ/mol) ± SD |
|---|---|---|---|---|---|---|
| ***Bc*WH2_CBM[b]** | CGM | 67 (± 6) | 1[d] | -24 (± 0.3) | -59 (± 11) | 35 (± 12) |
| | KGM | 52 (± 7) | 1[d] | -25 (± 0.3) | -56 (± 4) | 32 (± 4) |
| | M6 | 132 (± 34) | 1.3 (± 0.3) | -22 (± 0.7) | -47 (± 18) | 25 (± 18) |
| | M5 | 235 (± 3) | 1 (± 0.3) | -21 ± (0.1) | -68 (± 23) | 48 (± 23) |
| | M4 | 445 (± 21) | 1.7 (± 0.1) | -19 (± 0.1) | 32 (± 3) | 13 (± 3) |
| ***Bu*_CBM[c]** | CGM | 38 (± 6) | 1[d] | -6 (± 0.1) | -12 (± 1) | 6 (± 0.4) |
| | KGM | 47 (± 21) | 1[d] | -25 (± 1) | -54 (± 5) | 29 (± 6) |
| | M6 | 43 (± 2) | 0.8 (± 0.1) | -25 (± 2) | -64 (± 1) | 30 (± 1) |
| | M5 | 82 (± 6) | 1.3 (± 0.4) | -24 (± 0.2) | -39 (± 12) | 16 (± 12) |
| | M4 | 335 (± 26) | 0.7 (± 0.4) | -20 (± 0.2) | -117 (± 51) | 97 (± 51) |

[a] Parameters determined by ITC. Standard deviation calculated for the means of at least triplicate titrations.
[b] *Bc*WH2_GH26 isolated CBM.
[c] *Bu*-GH26 isolated CBM.
[d] Number of binding sites on CBM (N) was assumed as 1 for polysaccharides whose exact molar weight is unknown.

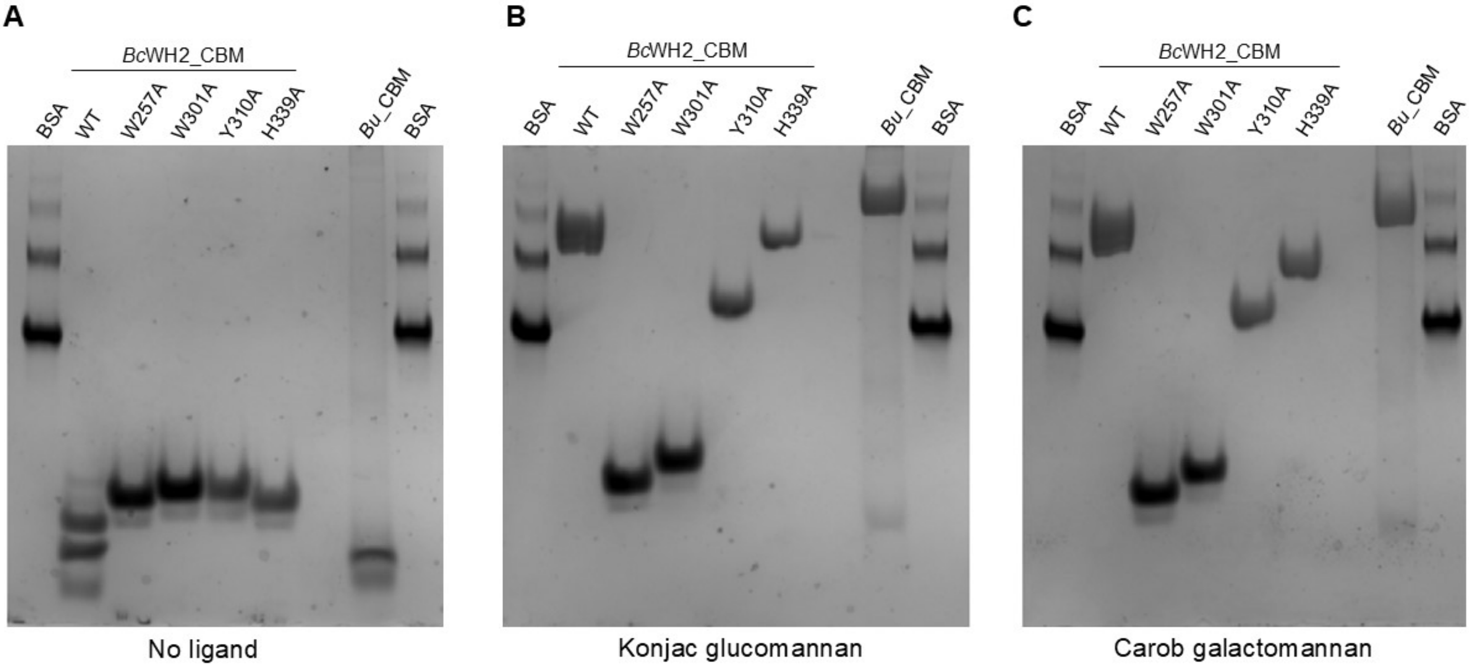


**Figure S5. Affinity gel electrophoresis of WT *Bc*WH2_CBM and *Bu*_CBM and mutants of *Bc*WH2_CBM against soluble mannans.** A) Negative control gel with no ligand added. B) Gel containing 1 mg/ml final concentration of konjac glucomannan. C) Gel containing 1 mg/ml final concentration of carob galactomannan. BSA was added in both the first and last lane of each gel as a non-polysaccharide-binding control.

**Table S8. Binding affinity of *Bc*WH2_CBM mutants against a variety of ligands as determined by ITC[a].**

| Construct | Ligand | $K_d$ (μM) ± SD | N ± SD |
|---|---|---|---|
| WT | Carob galactomannan | 67 (± 6) | 1[b] |
| | Konjac glucomannan | 52 (± 7) | 1[b] |
| | Mannohexaose | 132 (± 34) | 1.3 (± 0.3) |
| W257A | Carob galactomannan | NB[c] | - |
| | Konjac glucomannan | NB[c] | - |
| | Mannohexaose | NB[c] | - |
| W301A | Carob galactomannan | NB[c] | - |
| | Konjac glucomannan | NB[c] | - |
| | Mannohexaose | NB[c] | - |
| Y310A | Carob galactomannan | 254 (± 155) | 1[b] |
| | Konjac glucomannan | 142 (± 63) | 1[b] |
| | Mannohexaose | 348 (± 133) | 1.5 (± 0.6) |
| H339A | Carob galactomannan | 157 (± 88) | 1[b] |
| | Konjac glucomannan | 100 (± 15) | 1[b] |
| | Mannohexaose | 184 (± 17) | 0.67 (± 0.1) |

[a] Parameters determined by ITC. Standard deviation calculated for the means of at least triplicate titrations.
[b] Number of binding sites on CBM (N) was assumed as 1 for polysaccharides whose exact molar weight is unknown.
[c] No binding detected.

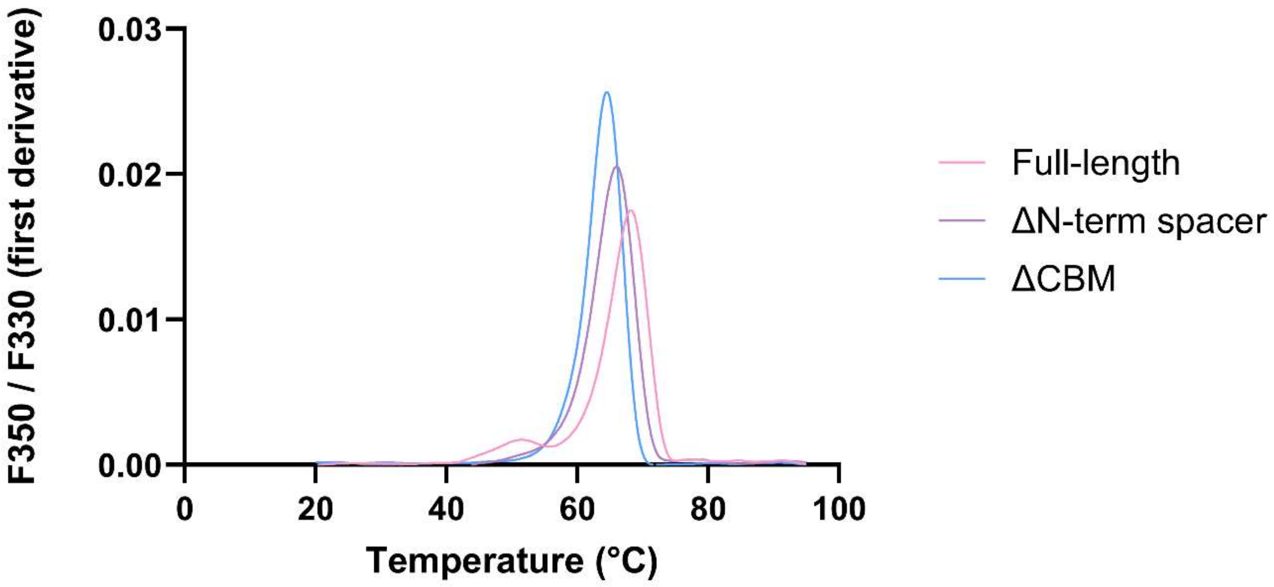


**Figure S6. Differences in thermostability between *Bc*WH2_GH26 and its truncated constructs determined using nanoDSF.** All three proteins were Thermal denaturant gradient between 20°C and 95°C was employed. The increase in the $F_{350}/F_{330}$ ratio indicates protein unfolding with the Tm indicated by the inflection point. The first derivative view was used to visualise the inflection point as the peak of the graph.

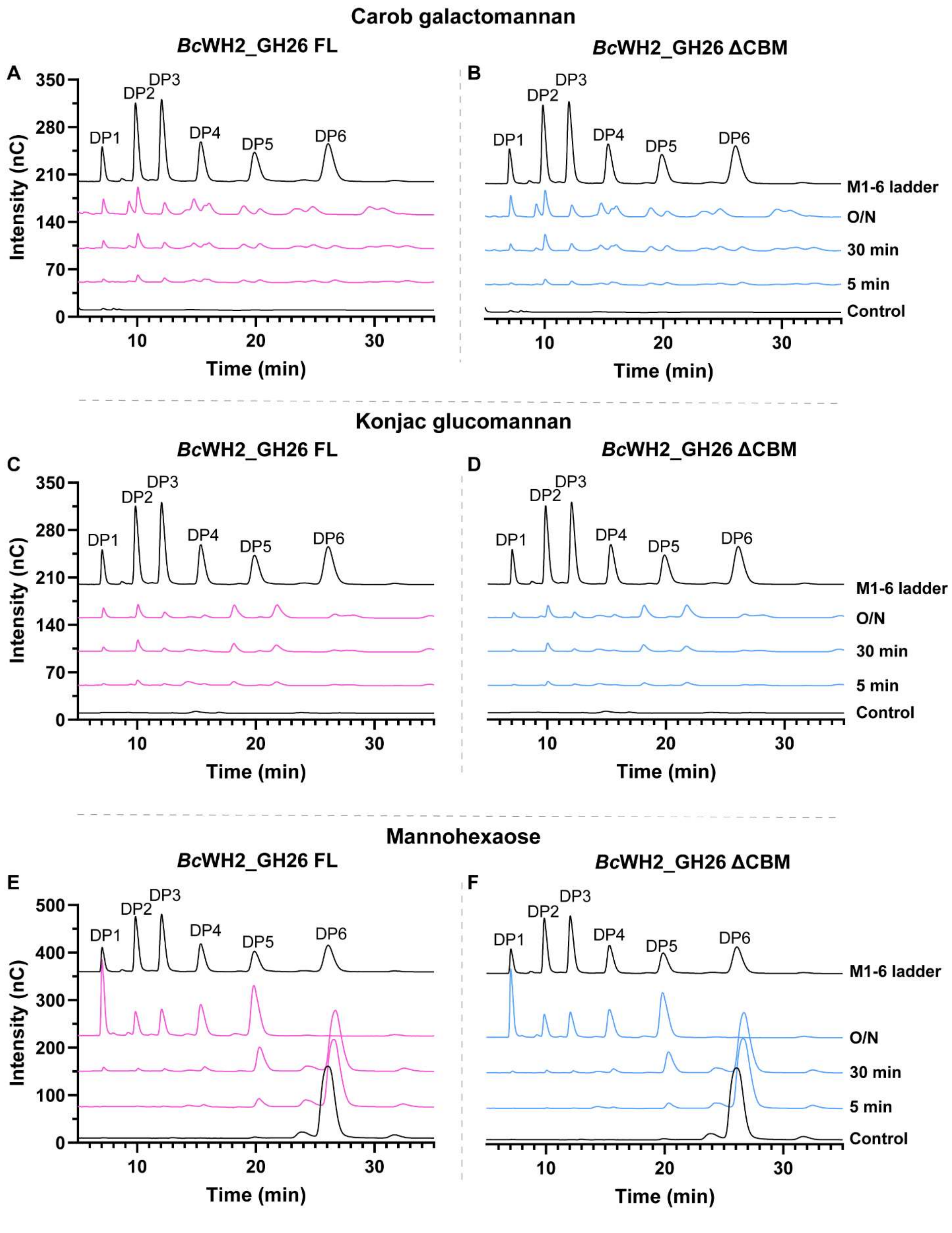


**Figure S7. Analysis of breakdown products of *Bc*WH2_GH26 endo mannanase vs mannans by ion-chromatography.** Carob galactomannan (A, B), konjac glucomannan (C, D) and mannohexaose (E, F). Hydrolysis products of *Bc*WH2_GH26 FL are shown in pink (left), and ΔCBM *Bc*WH2_GH26 in blue (right). The assays contained 1 μM final enzyme concentration; either 5 mg/ml final concentration of CGM or KGM, or 5 mM of mannohexaose; and 50 mM potassium phosphate buffer, pH7.5. The assays were incubated overnight (O/N) at 37°C, with samples taken at 5 and 30 minutes. Mannooligosaccharides (M1–M6) ladder is indicated in black as the top chromatogram, meanwhile the no enzyme control is shown in black as the bottom chromatogram.

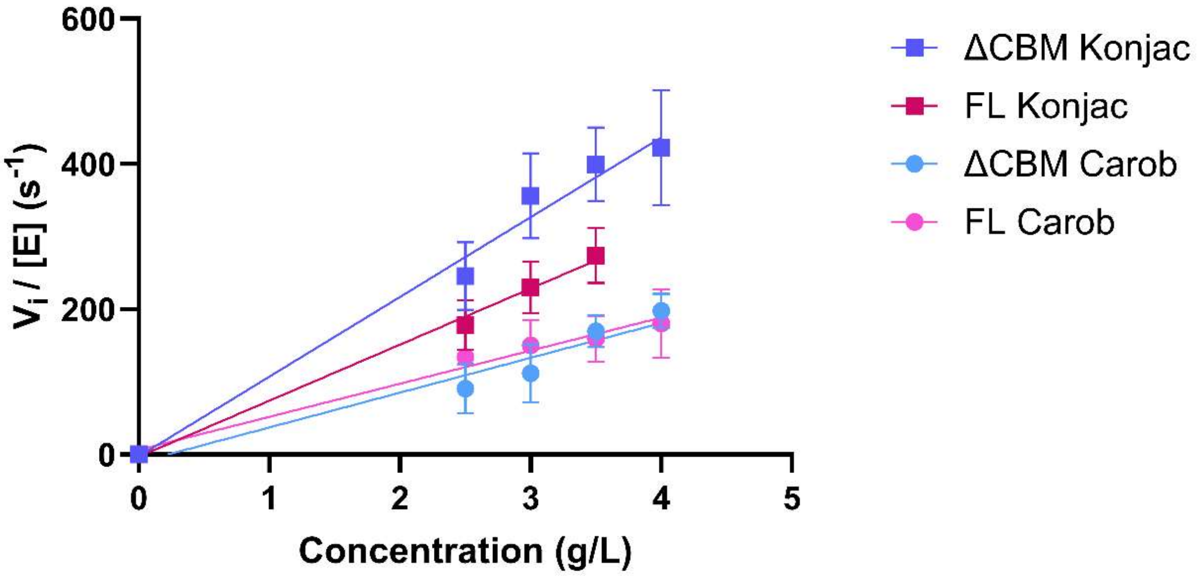


**Figure S8.** Linear regression used to estimate $k_{cat}/K_m$ values for *Bc*WH2_GH26 FL and ΔCBM *Bc*WH2_GH26 on konjac glucomannan and carob galactomannan.

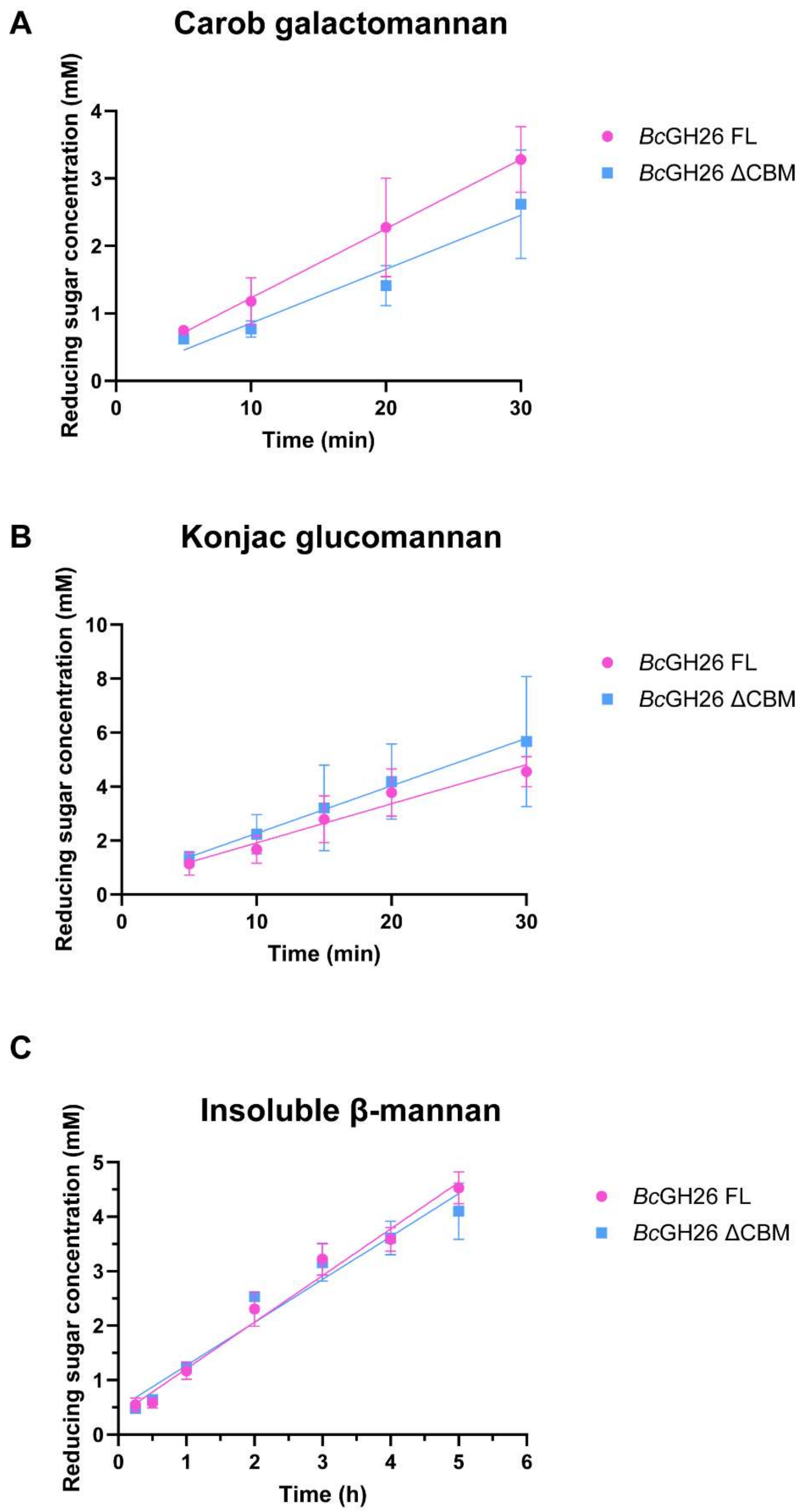


**Figure S9. Comparison of specific activity between full-length (FL) *Bc*WH2_GH26 and its truncated construct, ΔCBM *Bc*WH2_GH26.** For the soluble substrates, reactions contained 3 mg/ml final concentration of either A) carob galactomannan or B) konjac glucomannan; 10 nm final concentration of enzyme; and 50 mM of potassium phosphate buffer, pH 7.5. The assays were incubated at 37°C with 900 rpm agitation, with samples being taken at 5, 10, 20 and 30 min. For the insoluble substrate (C), the reactions contained 3 mg/ml insoluble mannan, 1 μM final enzyme concentration, and 50 mM of potassium phosphate buffer, pH 7.5. They were at incubated at 37°C with 1500 rpm agitation, with samples being taken at 15, 30 min, and then every hour for 5 hours. All the reactions were stopped by adding DNSA and boiling the samples for 10 minutes, before reading Abs at 540 nm and reducing sugar released quantified using a mannose std curve.

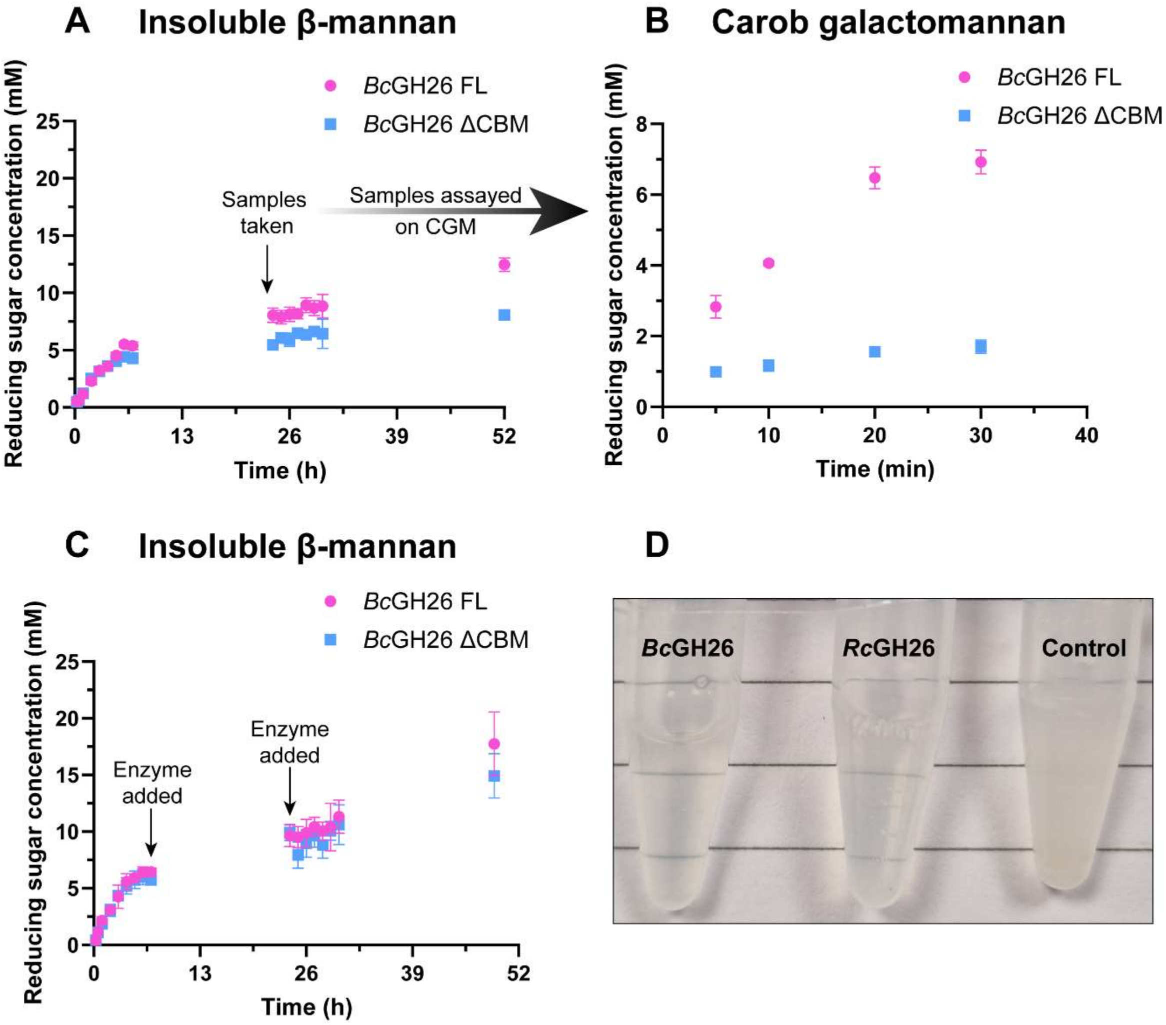


**Figure 10. Analysis of BcWH2_GH26 activity against insoluble β-mannan.** All the reactions contained 50 mM of potassium phosphate buffer, pH 7.5 and were incubated at 37°C with 1500 rpm agitation. A) Initial reactions contained 3 mg/ml of insoluble β-mannan and 1 μM of either FL *Bc*WH2_GH26 (in pink) or ΔCBM *Bc*WH2_GH26 (in blue), with samples being taken at various timepoints, then stopping the reactions by adding DNSA and boiling the samples for 10 minutes, before reading Abs at 540 nm. Reducing sugar released was quantified using a mannose std curve. After 24 h incubation time, an additional 40 μl sample was taken and assayed against carob galactomannan. B) Results of an assay of the taken samples (100 nM final enzyme concentration) against 5 mg/ml carob galactomannan. Samples were taken at 5, 10, 20, and 30 min. C) Repeat of the initial reaction shown in panel (A) with a modification involving addition of 1 μM of fresh enzyme aliquot at 7 h and 24 h incubation timepoints. D) Complete substrate solubilisation following 48 h incubation of the *Bc*WH2_GH26 and *Rc*GH26 with insoluble mannan. No enzyme control is shown on the right.

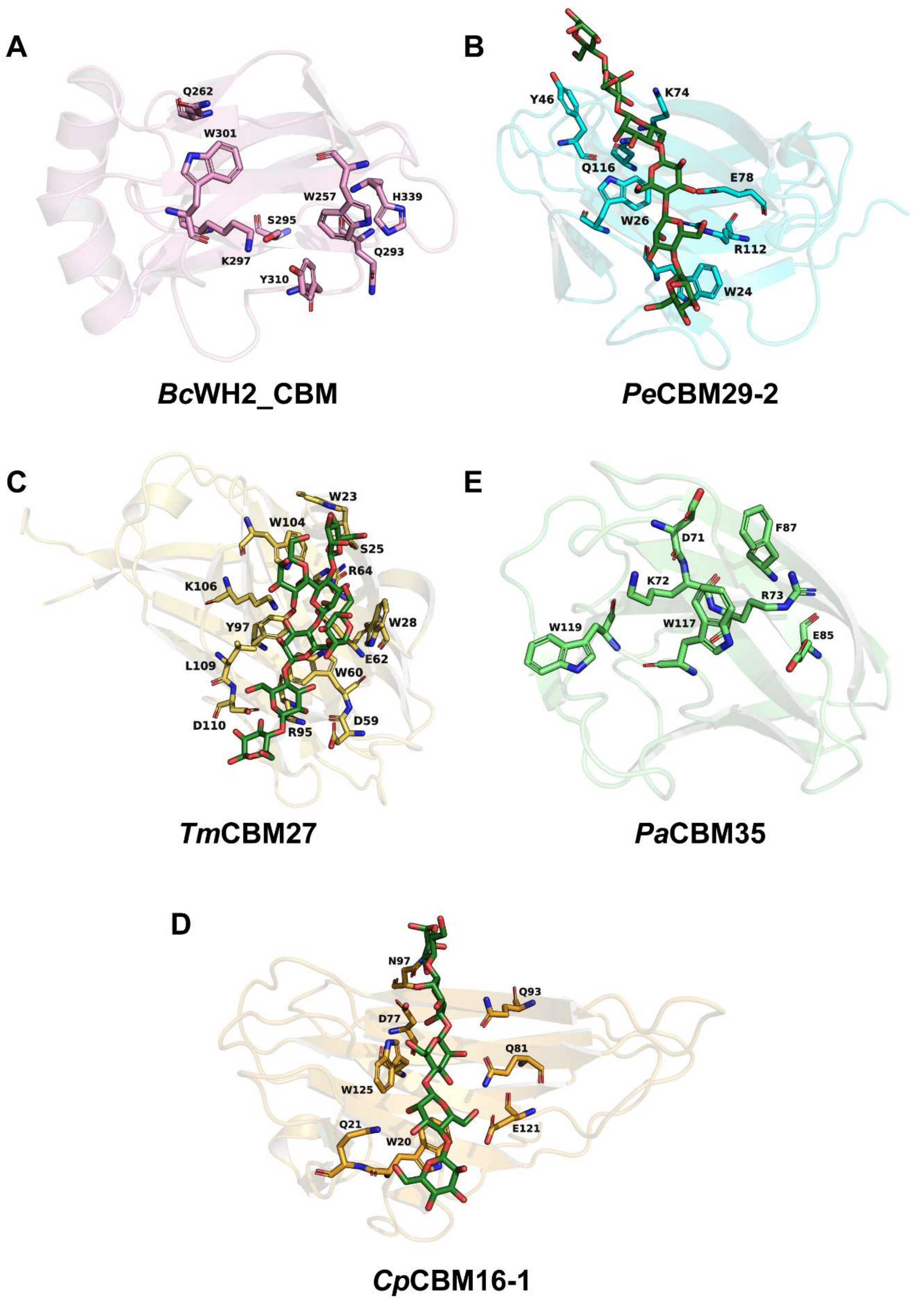


**Figure S11. Comparison of binding sites of selected β-mannan binding CBM families**. A) AlphaFold2 prediction of *Bc*WH2_CBM. B) Crystal structure of *Piromyces equi* CBM29-2 with M6 bound, PDB: 1GWL C) Crystal structure of *Thermotega maritima* CBM27 with G2M5 bound, PDB: 1OH4. D) Crystal structure of *Podospora anserina* CBM35, PDB: 3ZM8 E). Crystal structure of *Caldanaerobius polysaccharolyticus* CBM16-1 with M5 bound, PDB: 3OEB.
